# Solid-State Single-Molecule Sensing with the Electronic Life-detection Instrument for Enceladus/Europa (ELIE)

**DOI:** 10.1101/2022.08.31.505913

**Authors:** Christopher E. Carr, José L. Ramírez-Colón, Daniel Duzdevich, Sam Lee, Masateru Taniguchi, Takahito Ohshiro, Yuki Komoto, Jason M. Soderblom, M. T. Zuber

**Affiliations:** Daniel Guggenheim School of Aerospace Engineering, Georgia Institute of Technology, Atlanta, GA, 30332, USA; School of Earth and Atmospheric Sciences, Georgia Institute of Technology, Atlanta, GA 30332, USA; Massachusetts General Hospital, Department of Molecular Biology, Boston, MA 02114, USA; Howard Hughes Medical Institute, Boston, MA 02114, USA; Department of Chemistry, University of Chicago, 5735 S Ellis Avenue, Chicago, IL 60637; MIT Department of Electrical Engineering and Computer Science, Cambridge, MA 02139, USA; Osaka University, Institute of Scientific and Industrial Research, Osaka 565-0871 Japan; MIT Department of Earth, Atmospheric and Planetary Sciences, Cambridge, MA 02139, USA.

**Author notes:** Correspondence to. Address: ESM Building, Room G10, 620 Cherry St NW, Atlanta, GA 30332, USA. Co-first authors (Christopher E. Carr, José L. Ramírez-Colón).

**Keywords:** Life detection, Europa, Enceladus, Mars, biosignatures, amino acids, solid-state nanogaps, mechanically controlled break junction, HOMO–LUMO gap.

## Abstract

Growing evidence of the potential habitability of Ocean Worlds across our Solar System is motivating the advancement of technologies capable of detecting life as we know it – sharing a common ancestry or common physicochemical origin to life on Earth – or don’t know it, representing a distinct genesis event of life quite different than our one known example. Here, we propose the Electronic Life-detection Instrument for Enceladus/Europa (ELIE), a solid-state single-molecule instrument payload that aims to search for life based on the detection of amino acids and informational polymers (IPs) at the parts per billion to trillion level. As a first proof-of-principle in a laboratory environment, we demonstrate single-molecule detection of the amino acid L-proline at a 10 µM concentration in a compact system. Based on ELIE’s solid-state quantum electronic tunneling sensing mechanism, we further propose the quantum property of the HOMO–LUMO gap (energy difference between a molecule’s highest energy occupied molecular orbital and lowest energy unoccupied molecular orbital) as a novel approach to measure amino acid complexity. Finally, we assess the potential of ELIE to discriminate between abiotically and biotically derived (-amino acids in order to reduce false positive risk for life detection. Nanogap technology can also be applied to the detection of nucleobases and short sequences of IPs such as, but not limited to, RNA and DNA. Future missions may utilize ELIE to target preserved biosignatures on the surface of Mars, extant life in its deep subsurface, or life or its biosignatures in the plume, surface, or subsurface of ice moons such as Enceladus or Europa.

**One Sentence Summary:** A solid-state nanogap can determine the abundance distribution of amino acids, detect nucleic acids, and shows potential for detecting life as we know it and life as we don’t.

## 1. Introduction

Widespread synthesis of complex organics (amino acids, nucleobases, sugars) occurred early in the history of the solar nebula due to radiation processing of ices (Ciesla and Sandford 2012; Nuevo *et al*. 2012; Nuevo *et al*. 2009; Meinert *et al*. 2016), which is likely to occur throughout the universe. These building blocks of “life as we know it” are believed to have been delivered to all potentially habitable zones in the Solar System by comet and meteorite impacts (Schmitt-Kopplin *et al*. 2010; Engel and Macko 1997; Cooper *et al*. 2011; Martins *et al*. 2008; Callahan *et al*. 2011). Life might have arisen under similar physicochemical environments, such as within alkaline vent systems (Martin and Russell 2007; Martin *et al*. 2008) or impact-driven hydrothermal systems (Osinski *et al*. 2013) on the early Earth, Mars, or within subsurface oceans on icy worlds (Vance *et al*. 2007; Russell *et al*. 2014; Barge and White 2017). Because these building blocks form in interstellar space (Ciesla and Sandford 2012; Nuevo *et al*. 2012; Nuevo *et al*. 2009), in reducing planetary atmospheres (Hörst *et al*. 2012), and through aqueous chemistry (Ménez *et al*. 2018; Steel *et al*. 2017), we hypothesize that amino acids (AAs) are a common component of life, and that nucleic acids or related informational polymers (IPs) might be a common solution for information storage and heredity.

Amino acids can be stable over geologic time (Pizzarello and Cronin 2000; Glavin *et al*. 1999; Bada *et al*. 1998; Engel and Macko 1997), and the structural complexity and abundance distribution that has been measured from extraterrestrial material, as well as under terrestrial abiotic production, represents a null hypothesis for amino acid distributions in the search for ancient or extant life (Davila and McKay 2014; Reh *et al*. 2016). Quantum chemical calculations and biochemical experiments have further shown that the chemical reactivity of proteinogenic amino acids exhibits a notable pattern related to their temporal emergence in the genetic code (Granold *et al*. 2018); amino acids added later into the genetic code have greater reactivity than those found in the Murchison meteorite or recovered from the Miller-Urey experiment, which coincide with those that are predicted to have been added early (Trifonov 2009). Thus, as we discuss later, this property can be utilized as a new measure of amino acid complexity and possibly as an indicator of life that utilizes regulated electron transfer mechanisms. Therefore, the detection of amino acids, along with their abundance distribution, can be used as a potential biosignature that would allow for the differentiation between widespread abiotic chemistry and potentially ancient or extant life. Alternatively, charged backbones, like the polyphosphates of DNA and RNA, are likely universal for aqueous life (Benner 2017; Benner 2004): they separate a polymer’s physical properties from its associated information content, facilitating replication and evolution. Quantifying the presence of linear charged polymers with encoded information content could provide evidence of life, even without knowing the structure of the informational units (such as nucleobases) or their sequence. However, recent work demonstrating abiotic synthesis of RNA polymers 100-300 nucleotides in length on mineral glass surfaces demonstrates the potential for linear charged polymers in a prebiotic setting (Jerome *et al*. 2022).

Life elsewhere may also use similar building blocks because of shared ancestry through meteoritic exchange, a scenario most plausible for Earth and Mars (Carr 2022; Fritz *et al*. 2005; Shuster and Weiss 2005; Weiss *et al*. 2000; Gladman and Burns 1996a; Gladman *et al*. 1996b). Instruments targeting RNA or DNA, such as the Search for Extra-Terrestrial Genomes (SETG) (Ruvkun *et al*. 2002; Carr *et al*. 2016; Lui *et al*. 2011; Isenbarger *et al*. 2008a; Isenbarger *et al*. 2008b) are in development, but rely on biological reagents that must be stabilized, and require a complex series of steps to prepare nucleic acids for analysis. Solid-state nanopores have demonstrated the detection and characterization of nucleic acids, and have been proposed to potentially detect other polymers (Rezzonico 2014; Xia *et al*. 2022). However, at present, such nanopores require high (MHz) sample rates and ultra-low noise, and even so, solid-state nanopore detection of individual bases has not yet been realized (Shekar *et al*. 2016).

Here we propose the Electronic Life-detection Instrument for Enceladus/Europa (ELIE), a solid-state single-molecule detector. ELIE relies on quantum electronic tunneling (QET) nanogap sensors (Figure 1), which can detect and discriminate among single α-amino acids (Ohshiro *et al*. 2014), and detect RNA and DNA, including individual bases and, with caveats, short oligonucleotide sequences (Ohshiro *et al*. 2018; Ohshiro *et al*. 2012). We aim to primarily target amino acids and IPs, although we are developing ELIE with the potential to target other types of molecules, including abiotic or prebiotic polymers (Figure 2). This versatility of detecting a wide array of molecules would allow ELIE identify forward contamination, but also ancestrally related life – and possibly mirror-image life or alternative polymers. Here we focus on amino acid detection because amino acids exhibit relatively high stability under harsh conditions in comparison to IPs, and are potential targets for characterizing abiotic and prebiotic environments, and classifying ancient or extant life.

**Fig. 1.**
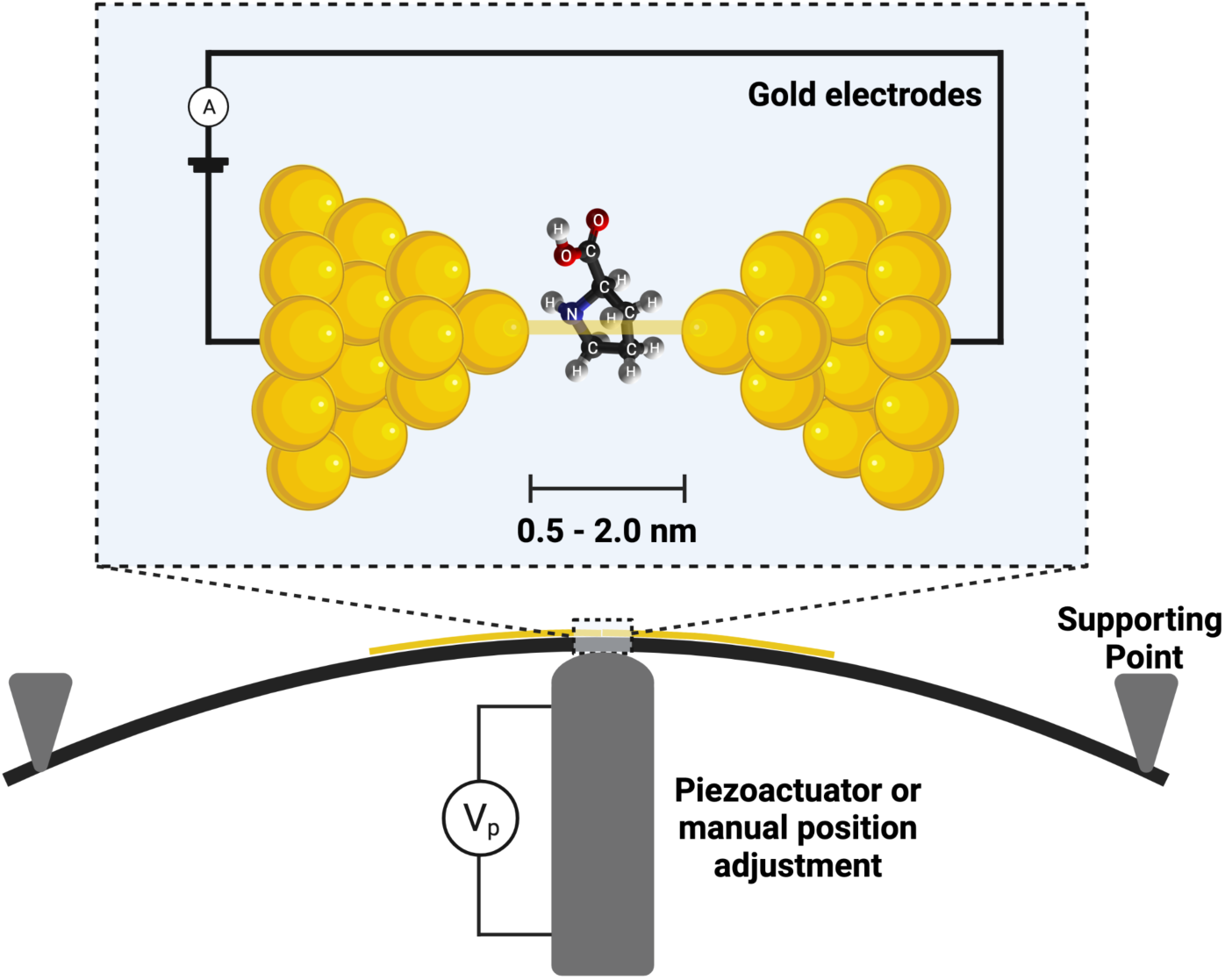
Single-molecule detection with a quantum electronic tunneling nanogap sensor, using the Mechanically Controlled Break Junction method. A gold nanowire is mechanically broken by three-point bending, forming two gold nanoelectrodes. Additional adjustments of the actuator or manual micrometer change the gap size. Due to the dependency of electron tunneling upon the electronic structure of any intervening molecule, target molecules are detected by variations in electrical current between the electrodes (Di Ventra and Taniguchi 2016).

**Fig. 2.**
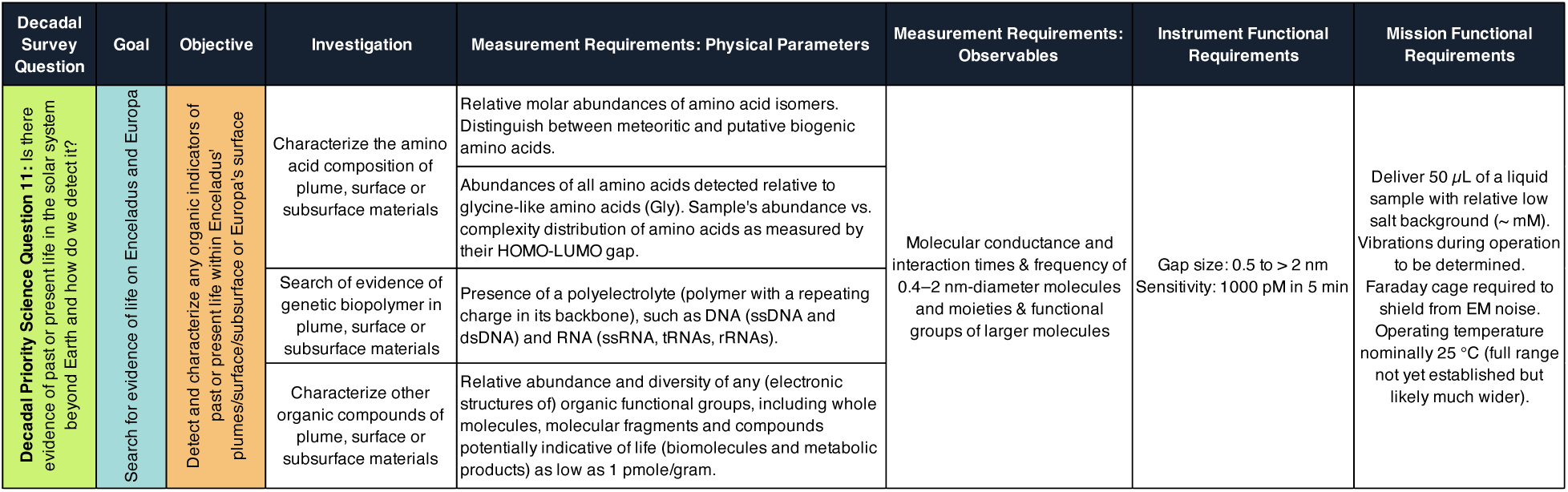
ELIE Science Traceability Matrix. Adapted for ELIE from goals of the Europa Lander Study Report and the Enceladus Orbilander Mission Concept (Hand *et al*. 2017; MacKenzie *et al*. 2021).

The discovery of Enceladus’ plume of icy grains and gas by the Cassini mission (Porco *et al*. 2006; Waite Jr *et al*. 2009) and the tentative evidence for transient water vapor plumes at several locations on Europa found in Hubble Space Telescope observations (Roth *et al*. 2014; Sparks *et al*. 2016), support the idea of material transport from the interior of these moons and, in concert with other studies, the presence of global subsurface oceans (Kivelson *et al*. 2000; Thomas *et al*. 2016; Zimmer *et al*. 2000). Based on known amino acids degradation rates, previous works suggest that a detection of amino acids in these predicted subsurface oceans above a concentration of 1 nM would indicate active production by geochemical or biotic pathways (Truong *et al*. 2019). Additionally, Steel *et al*. 2017 predict that methanogen-based microbial life and abiotic hydrothermal processes could produce a maximum of 90 µM and 104 µM concentrations of amino acids, respectively, in the predicted subsurface oceans of these moons.

In this study, we validate the single-molecule detection of L-proline with our first low-technology readiness level (TRL) ELIE prototype, and use this result to extrapolate an order-of-magnitude sensitivity of 1 nM. Furthermore, we provide evidence that the quantum property of the HOMO–LUMO gap describes the physical mechanisms underlying the conductance patterns of different proteinogenic amino acids obtained from the solid-state single-molecule detector.

Using amino acid abundance distributions reported in the literature, we then show the potential for ELIE to discriminate between abiotically and biotically derived α-amino acids. Finally, this work provides preliminary evidence for the feasibility of integrating ELIE into future *in-situ* life detection missions.

## 2. Methods

### 2.1. ELIE Instrument Hardware

The starting point for ELIE was a benchtop nanogap system developed by the Taniguchi laboratory at Osaka University as reported in Tsutsui *et al*. 2008a, Ohshiro *et al*. 2012, Ohshiro *et al*. 2014, and Ohshiro *et al*. 2018 (Figure 3a). The benchtop system consists of a picoammeter, a National Instruments (NI) computer, a custom Faraday cage enclosing stepping and piezo motors and a jig structure holding a nanogap chip, a piezo controller, and battery banks. One of the goals of the subsequently-developed ELIE prototype is to demonstrate a significant mass and volume reduction relative to this benchtop system. Compared to the benchtop system, the ELIE system consists of a low-noise amplifier for voltage supply, a laptop for data collection, and a Faraday cage holding an amplifier head stage, a nanogap chip, and a manual micrometer (Figures 3b and 3c). The nanogap chips (Figure 3d) are fabricated as previously reported by Tsutsui *et al*. 2008a. First, a silicon substrate is spin-coated with a thin polyimide layer. Gold nanojunctions (100 nm x 100 nm) are then patterned on top using a standard electron-beam lithography and lift- off technique. Subsequently, the polyimide beneath the junction is carved away by isotropic reactive ion etching, using O2/CF4 gas, to generate a free-standing gold nanowire bridge; this bridge is later formed into a nanometer-scale gap prior to measurement using the Mechanically Controlled Break Junction (MCBJ) method. An optional addition is a cover made of polydimethylsiloxane (PDMS), attached to the silicon substrate and treated with ozone plasma for bonding. This cover contains a microchannel that allows for the containment of a sample solution and its retention through reduced evaporation under room temperature conditions. The Faraday cage is connected to a low-noise and high-bandwidth amplifier (Chimera Instruments, VC100, 8 pA RMS at 100 kHz, with sampling up to 4 MHz) that supplies and controls the bias voltage (clamp ± 1V) and current measurements (±20 nA) of the nanogap.

**Fig. 3.**
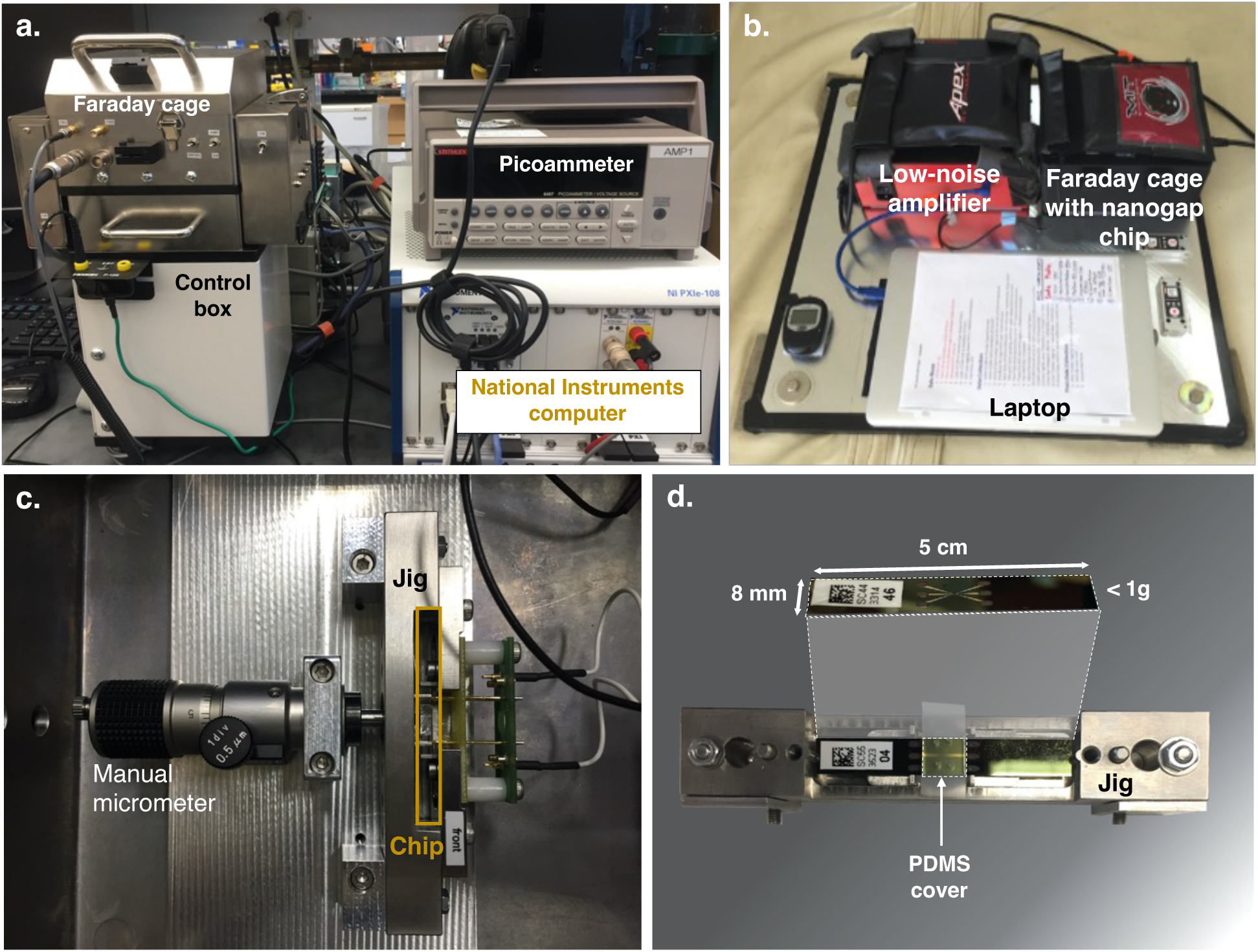
ELIE Instrument Prototype. **a**, Benchtop Osaka University Nanogap System used for the analysis of amino acids in Ohshiro *et al*. 2014. **b**, Current system consisting of a laptop controlling the low-noise amplifier that, in turn, supplies and controls voltage to and within a Faraday cage that encloses **c,** a manual micrometer and jig structure holding the **d,** nanogap chip.

### 2.2. ELIE Instrument Operation

The chip is mounted in a three-point bending mechanism in a jig within the Faraday cage, rinsed with 10% ethanol (as a wetting agent) and bent mechanically with a manual micrometer to form an atomically sharp gap. A 10 kΩ resistor is connected in series to protect the nanogap junction from over-current breakdown due to ohmic heating while in the connected state. In order to form the nanogap, the chip is bent by the micrometer to an opened state while monitoring the current. Once the gap is formed and in the open (unconnected) state, the resistor is removed from the circuit so that the current resulting from the applied voltage difference will reflect the gap conductance. As described by Tsutsui *et al*. 2008b, the resulting tunneling current (*I*) can be defined as an exponential function of the gap distance (d) as *I* ∼ exp (*β*d) with the decay constant 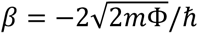, where *m* and *ħ* are the electron mass and the work function of Au. The gap is typically characterized in a dry state or in the presence of a buffer prior to making measurements with a sample. The distance is adjusted by means of chip bending, which produces changes in the gap distance that are a fraction of the vertical motion of the adjustment mechanism. The distance is monitored by the (VC100) measured current, which represents the sum of offset error (set to zero when bias is set to 0 mV), ionic current, and tunneling current. In practice, the baseline is zeroed during analysis so that differences in current reflect enhancement of tunneling current when a molecule of interest occupies the gap.

### 2.3. ELIE Instrument L-proline Measurement

L-proline was obtained from Sigma (81709) and a 10 µM L-proline solution prepared by dilution in nuclease free water with a background of 1 mM phosphate buffer, pH 7.4 at 25 °C (Sigma P3619). The solution was introduced through the microchannel and the current recorded for 5 minutes. Current measurements were conducted with an applied voltage of 100 mV and sampling rate of 4 MHz.

### 2.4. ELIE Instrument Tunneling Current Data Analysis

Data were low pass filtered to 25 kHz and downsampled to 100 kS/s in order to reduce the amount of electrical noise and the amount of data storage, respectively. Baseline adjustment was defined by estimating a kernel density over windows of 0.1 s with a 0.001 s sliding step. Events were detected using OpenNanopore v1.2, a tool that employs adaptive thresholding (by adjusting to low-frequency variations in the baseline) and a cumulative sum (CUSUM) fit to detect events (Raillon *et al*. 2012). Low-level noise signals within the set of events were subsequently identified using a 6σ threshold below the inferred mean of the events tunneling currents, and were excluded from further analysis. Features extracted from each of the events include the tunneling current (mean current level within single event) and dwell time (duration) (Figure 4a).

**Fig. 4.**
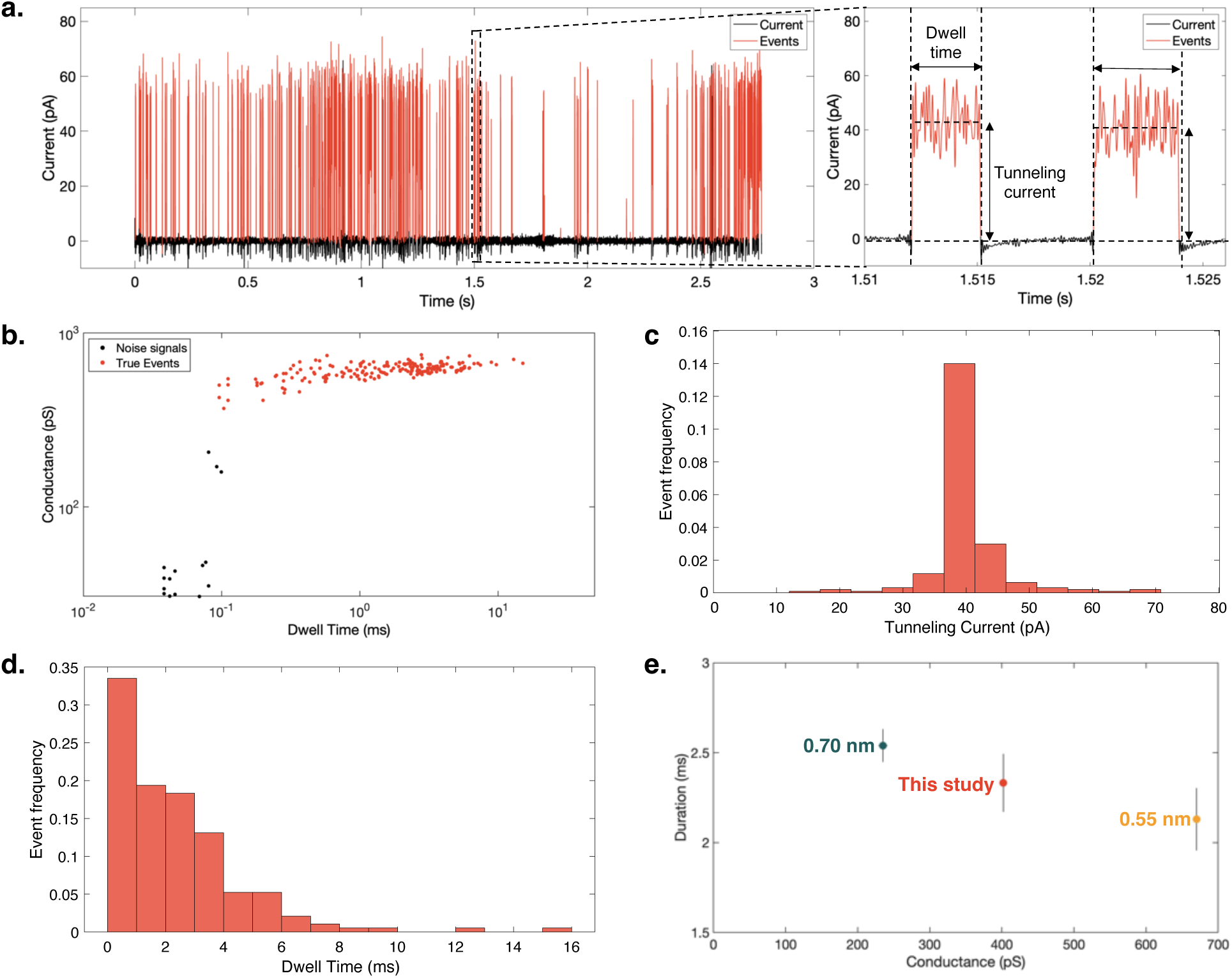
Single-molecule identification of L-proline. **a**, Conductance–time profile of 10 µM L-proline aqueous solution with an inset on the right showcasing the extracted features (dwell time and tunneling current) from each single event. **b**, Plot of single-molecule conductance and duration time of L-proline events displaying outliers within the data, identified by applying a 6σ threshold below the inferred mean of the event tunneling currents. **c**,**d,** Conductance and dwell histograms of L-proline events. **e**, Plot of single-molecule mean conductance and mean duration time of L-proline events obtained from 0.7-nm- and 0.55-nm-nanogap-electrodes measurements in Ohshiro *et al*. 2014 and by manually adjusting ELIE’s chip nanogap size.

### 2.5. Linear Regression Models of Amino Acid Conductance and Dwell Time

To gain insight into determinants of tunneling current and dwell time, linear regression models were fitted by plotting the values of various physical and quantum properties of proteinogenic amino acids (molecular weight, molecular volume, and HOMO–LUMO gap) against the conductance and dwell time values of amino acids measured by Ohshiro *et al*. 2014 (using the 0.55-nm and 0.70-nm-nanogap electrodes; refer to Figure S1). The HOMO–LUMO gap values of the proteinogenic AAs were reported by Granold *et al*. 2018 and determined by semiempirical calculations using the Molecular Orbital PACkage (MOPAC) 2003 AM1 Hamiltonian method in the MOPAC interface of Chem3D 9.0.

### 2.6. Predicting Differences in Distribution Patterns Between Meteoritic and Biologically Produced Amino Acids

We generated a literature-based database of reported abundances of meteoritic α-amino acids and amino acids extracted from diverse environmental samples (Elsila *et al*. 2021; Glavin *et al*. 2021; Aerts *et al*. 2020; Aponte *et al*. 2020; Glavin *et al*. 2020; Noell *et al*. 2018; Martins *et al*. 2015; Fuchida *et al*. 2014; Glavin *et al*. 2010; Hou *et al*. 2009; Martins *et al*. 2007). The analysis only considered meteoritic α-amino acids observed in multiple meteorites samples and reported by multiple authors. The studied meteorites were grouped into groups and subgroups, following mineralogy and elemental and isotopic composition (Table S1). Environmental samples considered in the analysis cover some of the closest terrestrial analogs to the surface of Mars and predicted subsurface environments of Enceladus and Europa (Table S2). Estimated event rates that would be expected for each amino acid by source (environmental or meteoritic) were determined by using an approximate standard event rate observed from the ELIE data in Section 2.4 of 50 events/s in 10 µM aqueous solutions, and assuming a linear relationship between concentration and event rate. Finally, single-molecule conductance values of α-amino acids reported by Ohshiro *et al*. 2014 (using the 0.70-nm-nanogap electrodes) were plotted against the predicted event rates. For the α-amino acids not reported by Ohshiro *et al*. 2014, the expected nanogap conductance signals were estimated using the HOMO–LUMO gap regression model obtained in Section 2.5.

## 3. Results

### 3.1. Proof of principle: Single-molecule detection of L-proline

Single-molecule detection was demonstrated with the ELIE system using 10 µM L-proline aqueous solutions and manual adjustment of the gap size between the electrodes. Manual adjustment limited maintaining a gap in the unconnected state within a measurable gap size range to short intervals on the scale of seconds. The current–time profile obtained from this experiment is shown in Figure 4a. After low-pass filtering, downsampling, and further processing through the OpenNanopore v1.2 software tool, 203 events were detected across the 2.7 s signal. By subsequently applying a 6σ threshold below the inferred mean of the event tunneling currents, 12 events were identified and removed, leaving 191 events to be analyzed (Figure 4b). As observed by the current and time histograms in Figure 4c and 4d, events detected had average current signals of 40.21 pA ± 6.12 pA and durations of 2.33 ms ± 2.21 ms. Extrapolation of this experimental result suggests a sensitivity of 1 nM or <100 ppt in 5 min, corresponding to 1 pmol/g. Lastly, the determined conductance was compared to measurements by Ohshiro *et al*. 2014 using the 0.55-nm and 0.70-nm-nanogap electrodes to test for L-proline.

Figure 4e suggests that the gap size was intermediate and closer to 0.70 nm. This results confirm the relationship between the gap distance and tunneling current previously outlined by Tsutsui *et al*. 2008b, where a larger gap distance generates lower tunneling currents, and therefore, conductance.

### 3.2. Linear regression models

A summary of the statistics obtained from the linear regression models, fitted by plotting conductance and dwell time values of amino acids against the specific amino acid’s molecular weight, molecular volume and HOMO–LUMO gap, is found in Table S3. The HOMO–LUMO gap showed the best linear relationship to the conductance and dwell time values obtained by the 0.55-nm and the 0.70-nm-nanogap electrodes of all tested covariates. However, only the linear relationship observed between the HOMO–LUMO gap and the conductance values of amino acids obtained from the 0.70-nm-nanogap electrodes is interpreted as statistically significant (Figure 5). Figures S2 and S3 showcase the remaining linear regression plots obtained from Section 2.5.

**Fig. 5.**
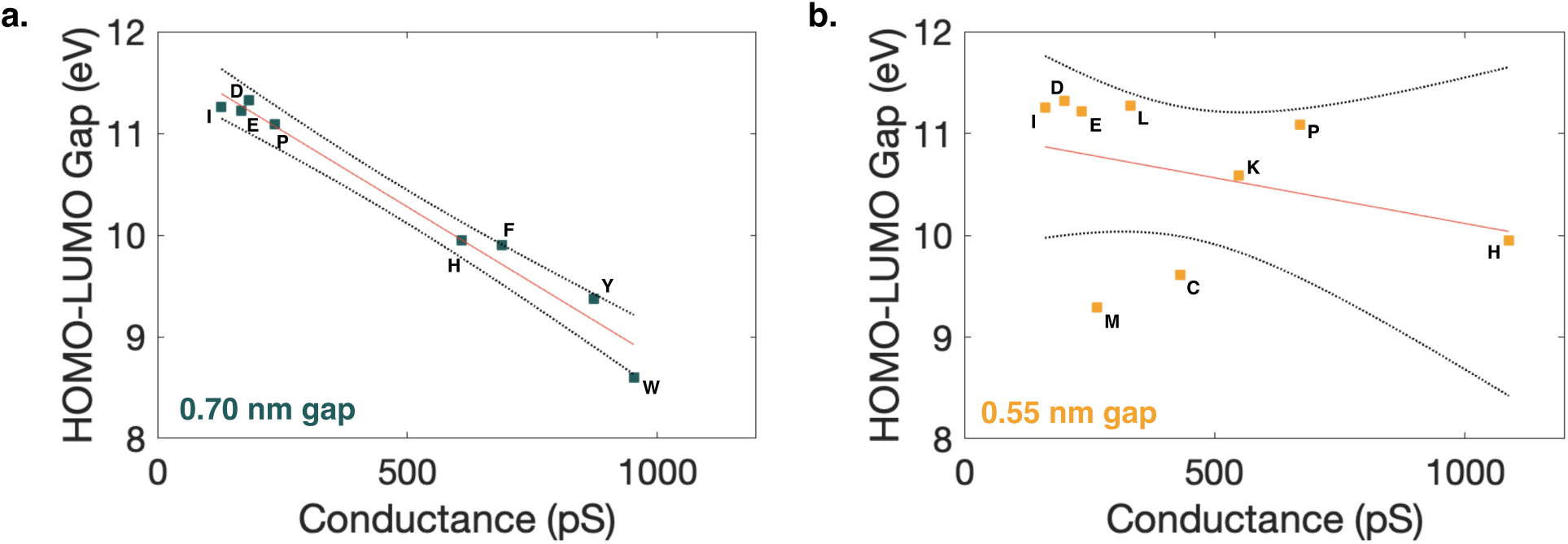
HOMO–LUMO gap of amino acids as a defining property for conductance levels using higher nanogap sizes. **a**,**b**, Linear regression plots of conductance measurements of proteinogenic amino acids, obtained by Ohshiro *et al*. 2014 using 0.70-nm- and 0.55-nm- nanogap electrodes, respectively, against their HOMO–LUMO gap values (as reported by Granold *et al*. 2018). Refer to Figures S2 and S3, and Table S3 for the remaining linear regression plots and their statistical summary. Refer to Dataset S1 for the key to the one-letter amino acid designations of the amino acids studied by Ohshiro *et al*. 2014.

### 3.3. Predicted event rates of biotically and abiotically derived α-amino acids

The conductance-event rate plots of amino acids by sample and category predicts event rates with orders of magnitude higher for amino acids derived from the environmental samples in comparison to the amino acids derived from meteoritic samples (Figure S4). Furthermore, an overlap of estimated event rates is observed between all the higher HOMO–LUMO gap amino acids in the genetic code (Figure 6), which also coincide with those α-amino acids found in meteorites, following the meteoritic reported amino acids gathered in the database.

**Fig. 6.**
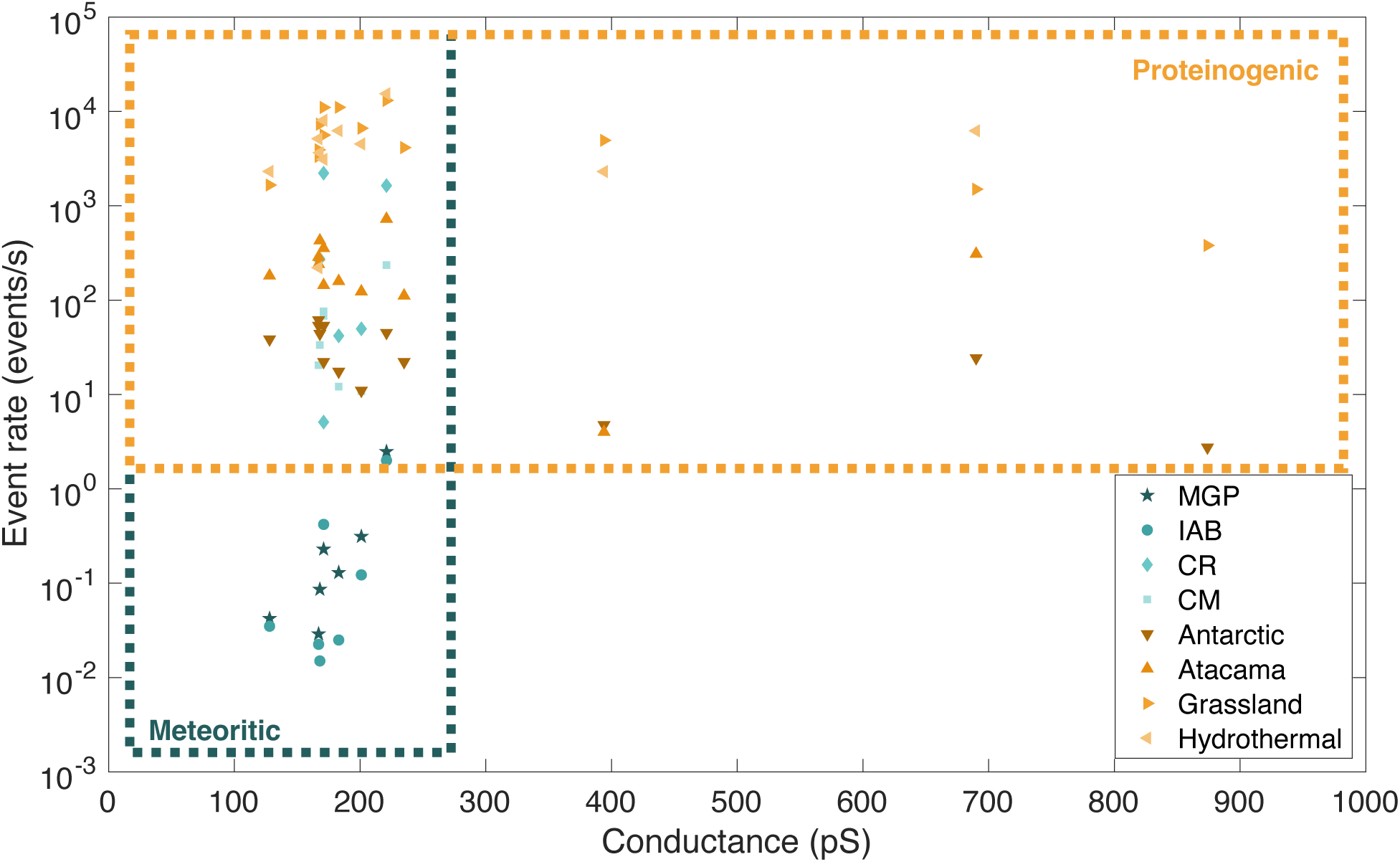
Potential discrimination between biotically and abiotically derived ɑ-amino acids. Distribution of predicted and experimentally validated conductance values of the 13 amino acids considered within the database as a function of the estimated event rates expected to be recorded from ELIE, on each of the meteorite (blue shades) and environmental (orange shades) samples. Meteorite group key: MGP (Main Group Pallasites); IAB (Iron meteorites group AB); CR (Renazzo-type chondrites); CM (Mighei-type chondrites).

## 4. Discussion

### 4.1. Nanogap technology for single-molecule detection and potential for space compatibility

Improving levels of detection towards single-molecule detection, in the pico-to femtomoles range, remains an ongoing goal for *in-situ* life-detection technologies. This is a significant issue because signatures of life or abiotically produced organics in other planetary bodies across our Solar System may be present at exceedingly low abundances, or only accessible in low abundances (Eigenbrode *et al*. 2021). Here, we demonstrate that the use of nanogap systems show great potential in this critical regard by performing as ultra-sensitive agnostic detectors without sacrificing the capacity to discriminate among molecules or chemical moieties (such as between different amino acids or DNA bases). Our extrapolated limit of detection meets thresholds proposed for organics in the Europa Lander notional Organic Compositional Analyzer baseline payload (Hand *et al*. 2017) and amino acids in the Enceladus Orbilander notional separation mass spectrometer and microcapillary electrophoresis with laser-induced fluorescence (MacKenzie *et al*. 2021). In comparison to other state-of-the-art instruments, such as gas chromatography-mass spectrometry (GC-MS) or capillary electrophoresis coupled to laser-induced fluorescence (CE-LIF), nanogap systems do not require derivatization agents to enable sensitive detection and separation of the targeted biomarkers.

Many derivatization agents have shown preferential reactions with water that produce increased background signals and low derivatization efficiencies of the targeted molecules (Goesmann *et al*. 2017; Leshin *et al*. 2013) and/or form by-products that interfere with the targeted molecules at low concentrations (Stalport *et al*. 2012; Creamer *et al*. 2017). Furthermore, the nanogap adjustment mechanism provides a general detection approach of nanometer-scale molecules (e.g., individual amino acids) or moieties (e.g., nucleobases) that are part of longer biomarkers, such as peptides, nucleic acids, or other polymers.

### 4.2. HOMO–LUMO gap as indicator of amino acid biogenicity

Linear regression models, obtained by using data of single-molecule conductance measurements from nanogap sensing experiments of single amino acids (Ohshiro *et al*. 2014), demonstrated a relationship between the level of conductance and the HOMO–LUMO gap values of 8 proteinogenic amino acids at higher nanogap sizes. Higher conductance values are obtained for amino acids with lower HOMO–LUMO gap values; a quantum property that defines the energy difference between a molecule’s highest energy occupied molecular orbital (HOMO) and lowest energy unoccupied molecular orbital (LUMO) in ground state. This may reflect the importance of electron transfer as an evolutionary adaptation of life as we know it; electron transfer pathways are essential for metabolism, biosynthesis, metal ligation, and counteracting biochemical changes in the environment, such as the rising oxygen concentrations within cells that initially evolved to thrive in anoxic environments (Cavalier-Smith 2006; Weiss *et al*. 2016; Moosmann 2021). Such potential molecular adaptations also correlate with various thermodynamic properties of amino acids within the genetic code: posited late additions of amino acids to the genetic code may have enabled increasingly reactive proteins to act as organic semiconductors (Figure 7), mediated by the incorporation of aromatic systems and less electronegative heteroatoms into their structures (Figure 8) (Bender *et al*. 2008; Gray and Winkler 2015).

**Fig. 7.**
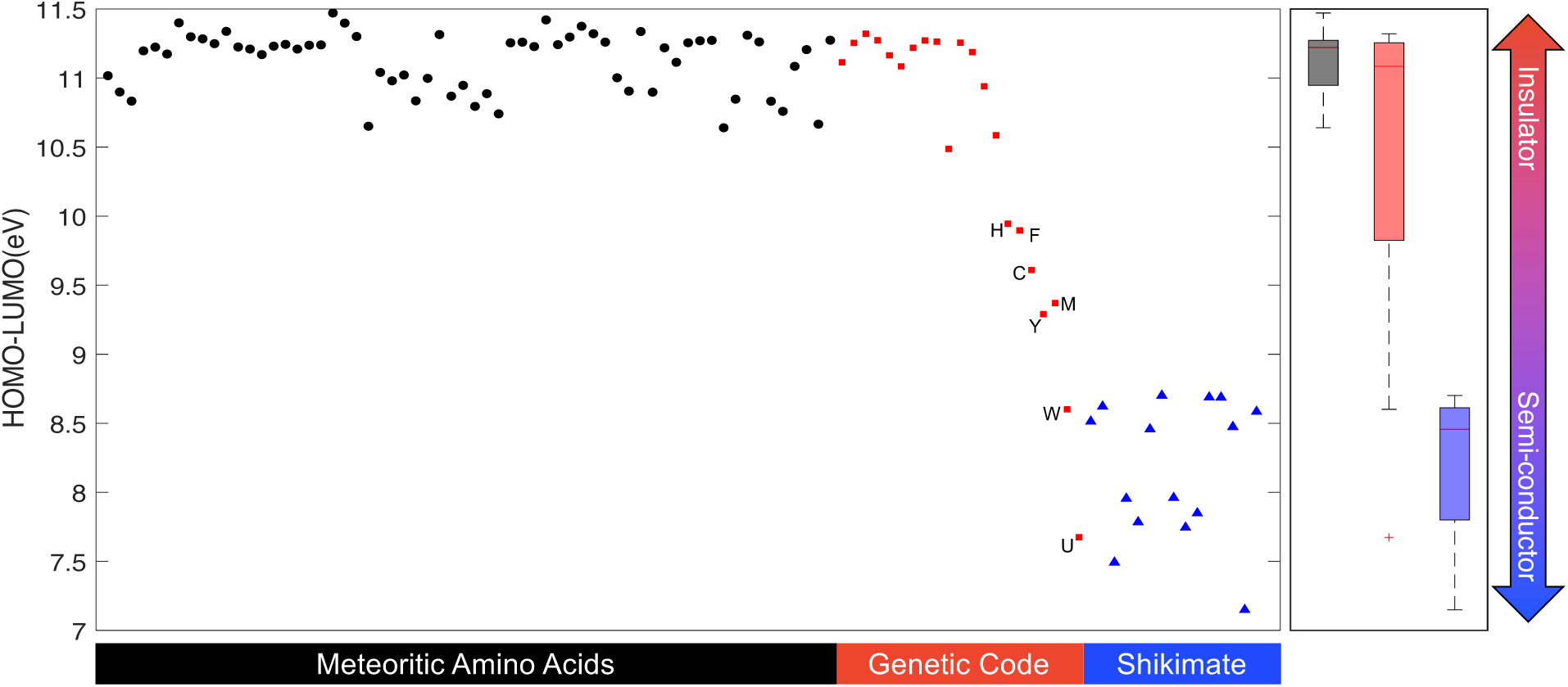
Amino Acid HOMO–LUMO gaps. Distributions of the HOMO–LUMO gaps of 62 Murchison meteorite amino acids, 21 genetically encoded amino acids, and various metabolic descendants of the shikimate pathway (a biosynthetic pathway that leads to the formation of F, Y and W). The quantum-chemical properties were determined by Granold *et al*. 2018 using semiempirical calculations with the MOPAC2003 AM1 software package. The 21 proteinogenic amino acids (including selenocysteine) are plotted in the consensus order of their evolutionary appearance according to Trifonov 2009. Late additions to the genetic code are labeled with their respective one-letter code (H: histidine; F: phenylalanine; C: cysteine; M: methionine; Y: selenocysteine). Refer to Dataset S2 for individual names and HOMO–LUMO gaps of each amino acid and metabolic descendant of the shikimate pathway.

**Fig. 8.**
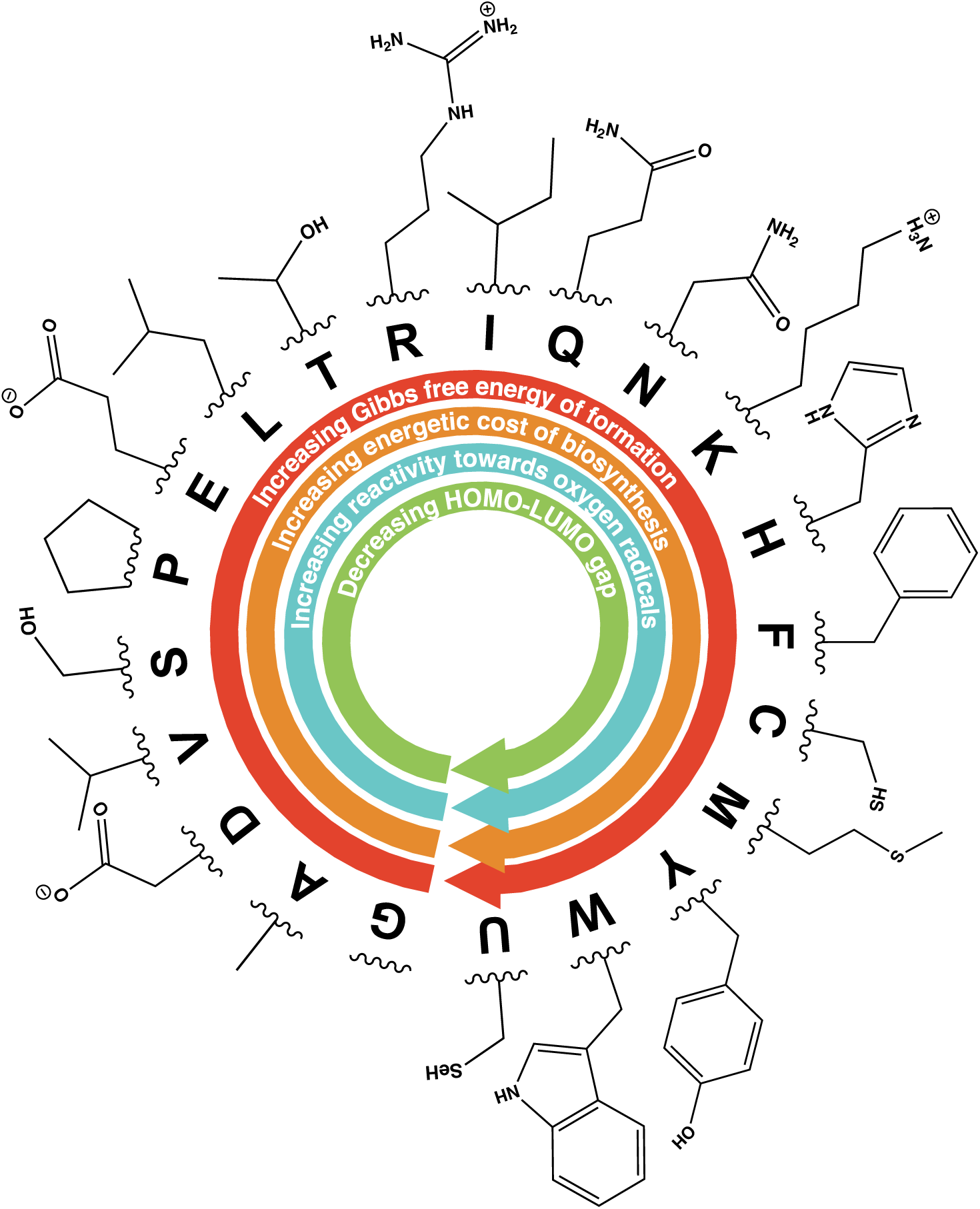
Consensus order of appearance of proteinogenic amino acids within the genetic code as predicted by Trifonov 2009, along with their functional groups. Arrows are showing increasing and decreasing patterns of some AAs quantum, redox, and thermodynamic properties as discussed and reported by Akashi and Gojobori 2002, Amend and Shock 1998, and Granold *et al*. 2018. Arrows indicate trends with quantitative support (Figure S7). One letter amino acid designations are provided in the Dataset 3.

From a quantum physical context, the relationship between the level of conductance and the HOMO–LUMO gap in solid-state single-molecule detectors is attributed to the electron transport mechanism behind single-molecule junctions. The band structures of the gold nanojunctions within the nanogap chip are composed of crystal orbitals that conduct electrons at the Fermi level, a continuous energy level at room temperature (Taniguchi, 2017). Once these orbitals interact with another molecule passing through the nanogap, electronic interactions occur, shifting electron transmission within the nanogap and, in turn, generating changes in the conductance band that are mainly dependent on the conformation of the molecule. Molecules with lower HOMO–LUMO gaps tend to have their HOMO orbitals overlapping to a greater extent with the Fermi level of the nanoelectrodes, thus permitting a greater electron transmission (Taniguchi 2019). Therefore, this quantum property of amino acids explains the trends observed in the resulting conductance of amino acids sensed by the nanogap system. Consequently, the HOMO–LUMO gap is an accurate predictor of amino acid conductance, and thus amino acid complexity, in nanogaps systems at nanogap sizes that are significantly higher that the targeted molecules (Figures 5). In contrast, this relationship is not true for mass and spatial size (molecular volume), which seem to be the determining mechanisms responsible of electrical sensing (via ionic current blockade) in biological and solid-state nanopores (Wei *et al*. 2020).

In contrast, no statistically significant relationship was observed between the dwell time from the nanogap sensing experiments of single amino acids by Ohshiro *et al*. 2014 and the proteinogenic amino acid’s physical and quantum properties considered in this study. However, as seen in Figure S5, there is an evident pattern in the dwell time at lower nanogap sizes, in which dwell time decreases in relation to the functional group of the amino acids tested. This can be related to the bonding strength between the gold nanoelectrodes and the anchoring group of the amino acid. It has been previously reported that thiol and amine groups form strong-stable and weak-covalent bonds on gold surfaces, respectively (Xue *et al*. 2014; Leff *et al*. 1996).

Furthermore, imidazole groups are known to coordinate to gold nanoparticles and contribute significantly to interactions between metallic surfaces and biological filaments (Zhou *et al*. 2010). These observations suggest that the bond strength between the gold atoms and amino acids like histidine, cysteine, and lysine allow for longer periods of single-molecular junctions at low nanogap sizes. Further studies are needed to assess other factors (e.g. electrical potential, molecular volume) and their possible combinatory influence on the dwell time at longer nanogap sizes. These results also highlight the need for automated gap control in order to maintain the gap at optimal distances for the molecular junction to be formed with the targeted molecules.

### 4.3. ELIE’s potential for discrimination between biotically and abiotically derived α-amino acids

Detection of low HOMO–LUMO gap small molecules may be indicative of life, but such molecules are not unique to living systems. Some amino acids in the genetic code with lower HOMO–LUMO gaps (<10 eV; 5σ below the mean HOMO–LUMO gap of Murchison meteorite amino acids, as reported by Granold *et al*. 2018) have been reported to be produced abiotically in terrestrial environments and endogenously in extraterrestrial materials. Ménez *et al*. 2018 demonstrated the abiotic formation of tryptophan, an aromatic α-amino acid with a low HOMO– LUMO gap, during hydrothermal alteration of mantle rock in the Atlantis Massif. Moreover, Friedel-Crafts-type reactions are predicted to be responsible for the formation of this amino acid during the hydrothermal alteration of oceanic peridotites; a different pathway from the Strecker- cyanohydrin synthesis, which has been predominantly supported for the formation of endogenous α-amino acids in CM and CR chondrites (Martins *et al*. 2007; Ehrenfreund and Charnley 2000; Glavin *et al*. 2020).

ELIE will be able to provide the abundance distributions of α-amino acids within the sample, and thereby assess the abiotic or biotic origin of these targeted molecules. Because the abiotic production of amino acids is thermodynamically driven (Higgs and Pudritz 2009), lower molecular weight amino acids, with lower values of Gibbs free energy of formation (Figure 8), are often found in high abundances within abiotic samples (Figure S6). Biotic production, on the other hand, is driven by functionality, yielding higher abundances of high molecular weight species with low HOMO–LUMO gaps (e.g. phenylalanine and tyrosine) relative to lower molecular weight amino acids. Therefore, if the mechanisms of formation mentioned above are ruled out, and the abundances of low HOMO–LUMO gap amino acids are high, then such detections (many events at high conductance) would be consistent with a potential biological source of amino acids or other small molecules.

### 4.4. Advancing ELIE

Future work includes evaluating ELIE’s potential for detection of biomolecular mixtures, non-standard nucleic acid bases and other polymers in order to expand the structural diversity scope of detection and reduce the risk of false negatives. This will allow the development of a biomolecule detection/characterization algorithm with the potential for molecule classification using supervised and unsupervised methods (Taniguchi *et al*. 2019). Furthermore, we seek to develop ELIE into a breadboard system with long-term expectations of being applicable to life detection missions in Ocean Worlds, satisfying the mass, power, and volume constraints needed for such missions. To reach these next steps, we are improving some of the instrument’s current limitations by (i.) integrating an automated piezoactuator for gap control, (ii.) optimizing the Mechanically Control Break Junction process, (iii.) characterizing gap quality through monitoring of quantized gap conductance, (iv.) automating sample handling, and (v.) reducing the size of the low-noise amplifier. These improvements will allow us to better assess the performance of ELIE in relevant environments, including the influence of salt, vacuum exposure, vibration tolerance, and effects of space radiation.

## 5. Conclusions

The discovery of a second genesis of life on Europa or Enceladus would immediately suggest life is common in the universe, addressing fundamental questions, about life’s origins and its distribution across the universe, that would change humanity’s perspective. The present efforts are a proof of concept of an instrument capable of detecting single amino acids, at a sensitivity that is not currently feasible for most of the emerging technologies for *in-situ* amino acid detection proposed for planetary missions to Ocean Worlds. The detection of the amino acid L-proline was validated in the µM range, and extrapolation suggests an ultimate sensitivity in the pM range. Furthermore, the quantum property of HOMO–LUMO gap is shown to best describe the conductance levels of proteinogenic amino acids obtained by previous works, and thus, proposed as a novel approach to measure amino acid complexity. Finally, by predicting event rates based on amino acid distributions in terrestrial and extraterrestrial samples, we suggest a potential approach to distinguish between biotically and abiotically derived amino acid distributions. Ongoing efforts aim to extend ELIE’s capabilities for detection of other biomolecules and to develop the instrument into a breadboard system for potential opportunities for infusion into Ocean Worlds missions.

## Acknowledgments

We thank Bernd Moosmann for providing us the HOMO–LUMO gap data from the Murchison meteorite amino acids, the genetically encoded amino acids, and the various metabolic descendants of the shikimate pathway and the data of the peroxyl radical scavenging assays.

## Author contributions

C.E.C. and J.L.R.C. conceived and wrote the paper; C.E.C. collected the data; C.E.C. and S.L. built the hardware; M.T., T.O. and Y.L. provided the nanogap sensors; C.E.C and D.D. designed the L-proline experiment; all authors edited and approved the manuscript.

## Author Disclosure Statement

The authors declare no competing interests.

## Funding

This work was supported by NASA awards 80NSSC19K1028 and 80NSSC22K0188 to C.E.C.

## Supplementary Data

**Figure S1.**
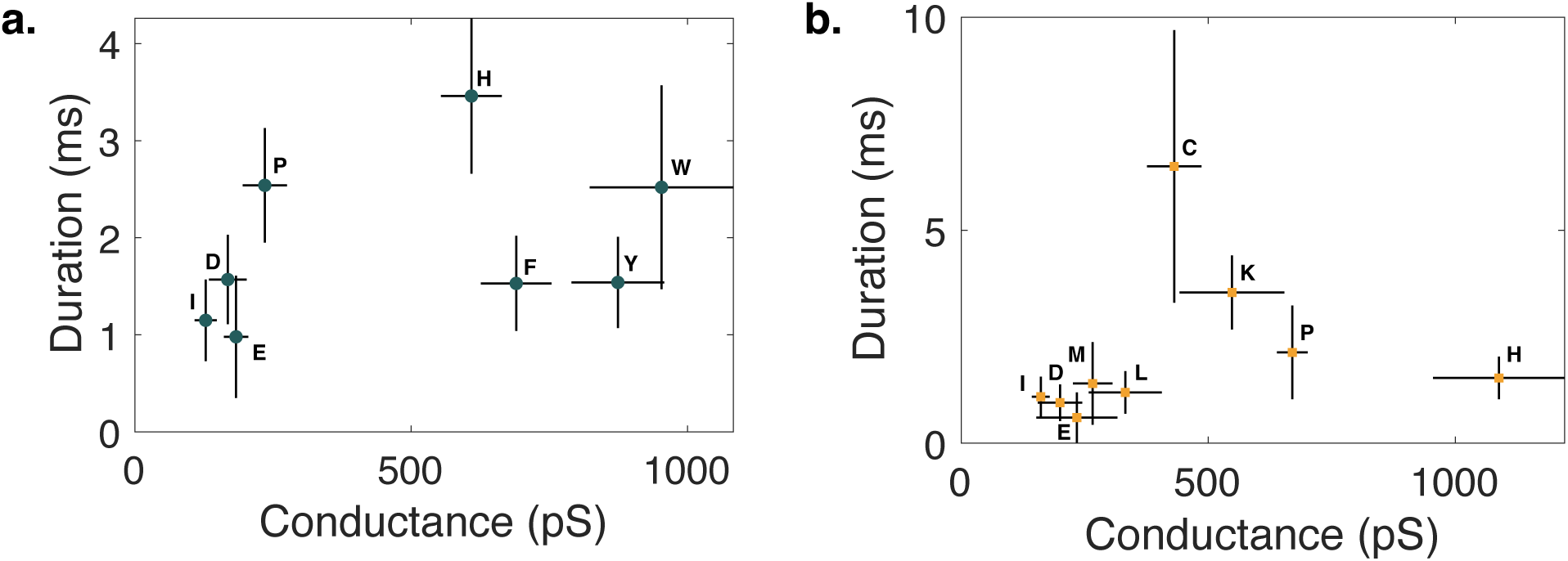
Two-dimensional plots of single-molecule conductance and duration time for 12 amino acids obtained by Ohshiro *et al*. 2014 using **a,** 0.7-nm- and **b,** 0.55-nm-nanogap electrodes. Conductance and duration times denote the peak conductance and peak duration times on their histograms. Error bars were calculated by σ/N^0.5^, where σ and N denote the standard deviation and total number of nanogap chips, respectively. Refer to Dataset S1 for the key to the one-letter amino acid designations and conductance and dwell time values.

**Figure S2.**
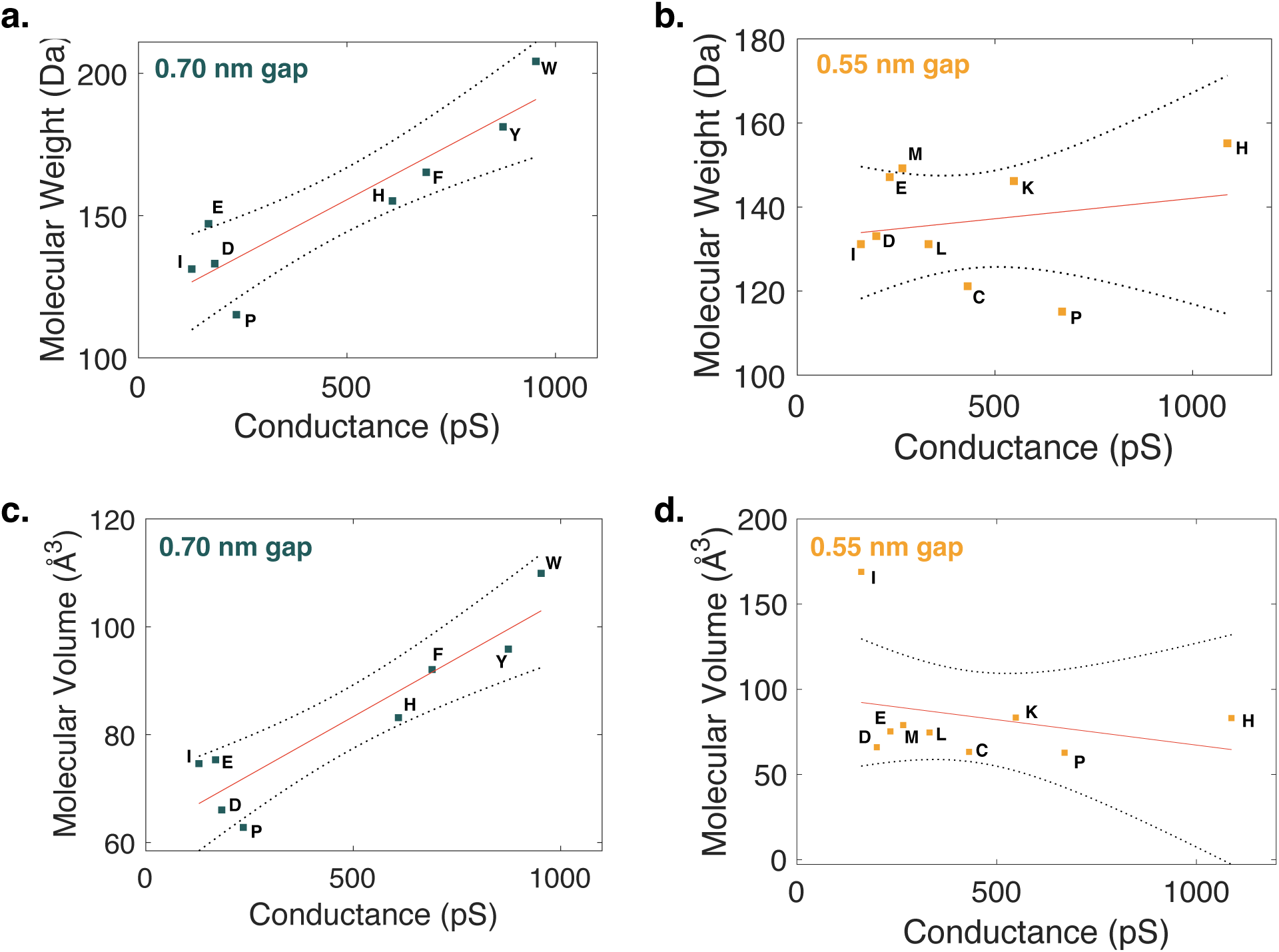
Linear regression plots of conductance measurements for 12 amino acids obtained by Ohshiro *et al*. 2014 using the **a,c,** 0.7-nm- and **b,d,** 0.55-nm-nanogap electrodes against their molecular weight and volume values.

**Figure S3.**
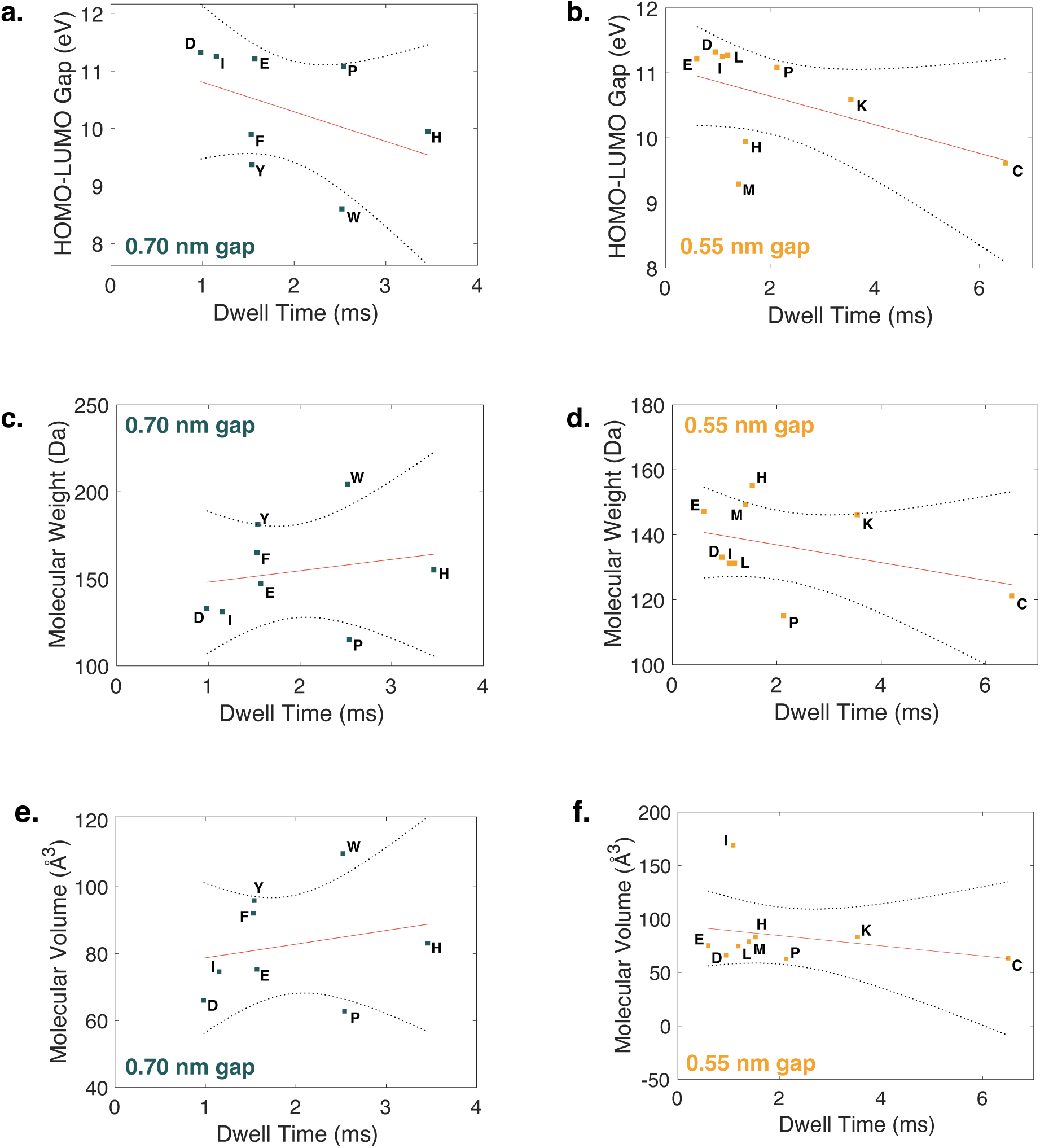
Linear regression plots of dwell time measurements for 12 amino acids obtained by Ohshiro *et al*. 2014. using the 0.7-nm- (blue dots) and 0.55-nm- (orange dots) nanogap electrodes against their **a,b,** HOMO-LUMO gap (as reported by Granold *et al*. 2018), **c,d,** molecular weight and **e,f,** volume values.

**Figure S4.**
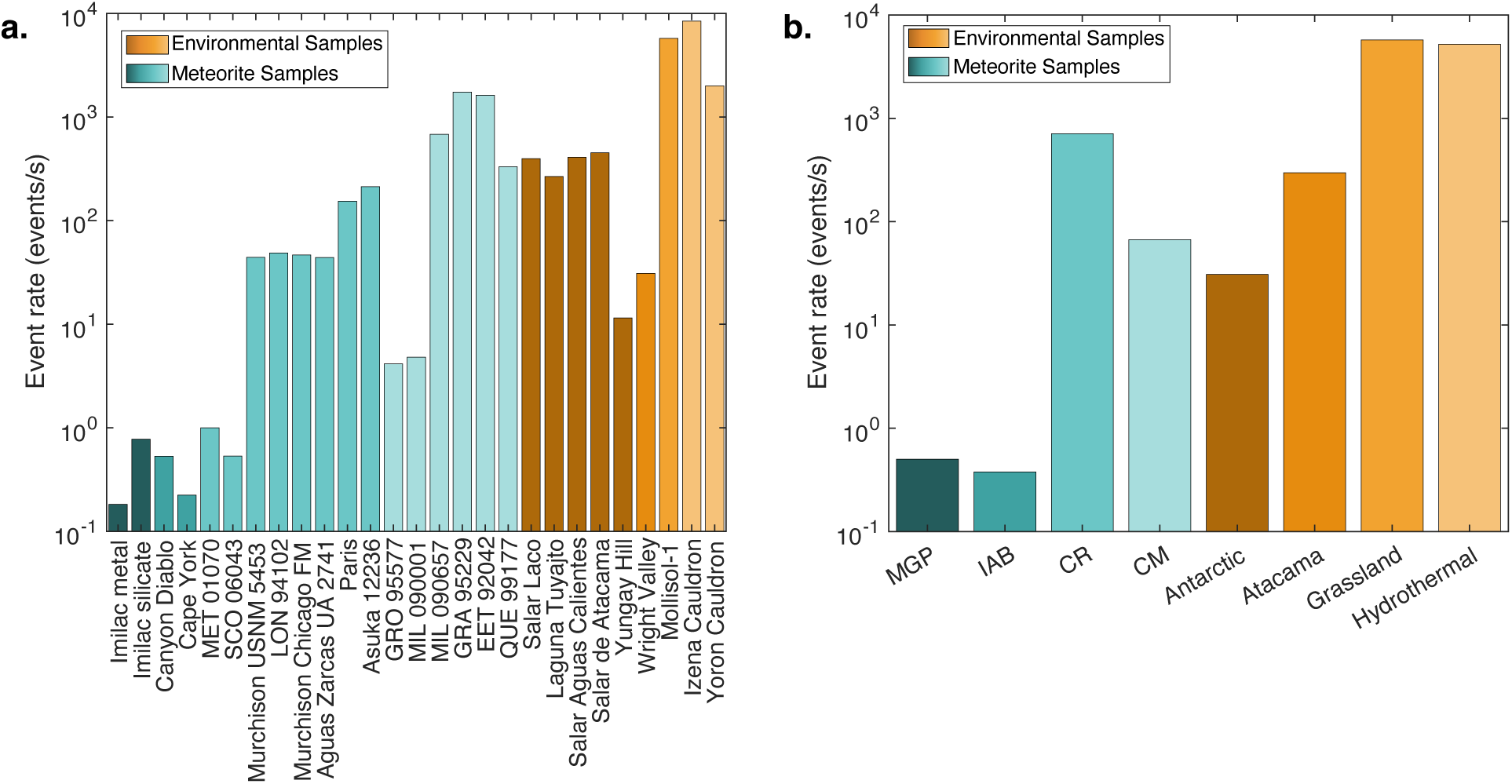
Predicted total event rates of ⍺-amino acids derived from biotic and abiotic sources. Classified by **a,** sample and **b,** category of the meteorite and environmental samples studied (tables S1 and S2). Such predicted events were determined by using an approximate standard event rate observed from the ELIE data of 50 L-proline events/s in 10 µM aqueous solutions, and assuming a linear relationship between concentration and event rate.

**Figure S5.**
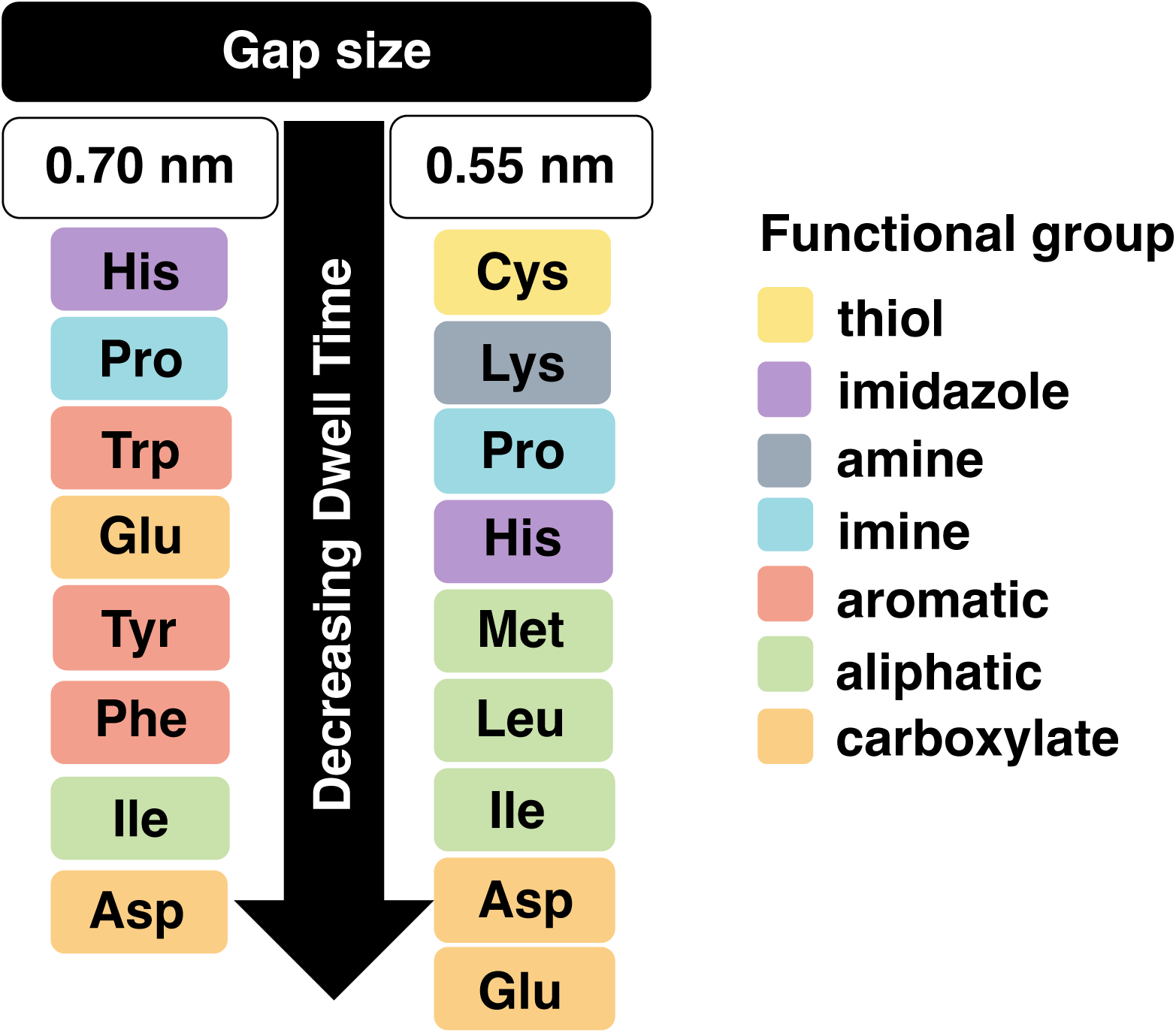
Dwell time and amino acid functional group correlation. Relationship between the single-molecule dwell times of the 12 amino acids obtained by Ohshiro *et al*. 2014 using 0.7- nm- and 0.55-nm-nanogap electrodes and their respective functional groups. Refer to Figure 8 for the chemical structure of the functional group pertaining each amino acid.

**Figure S6.**
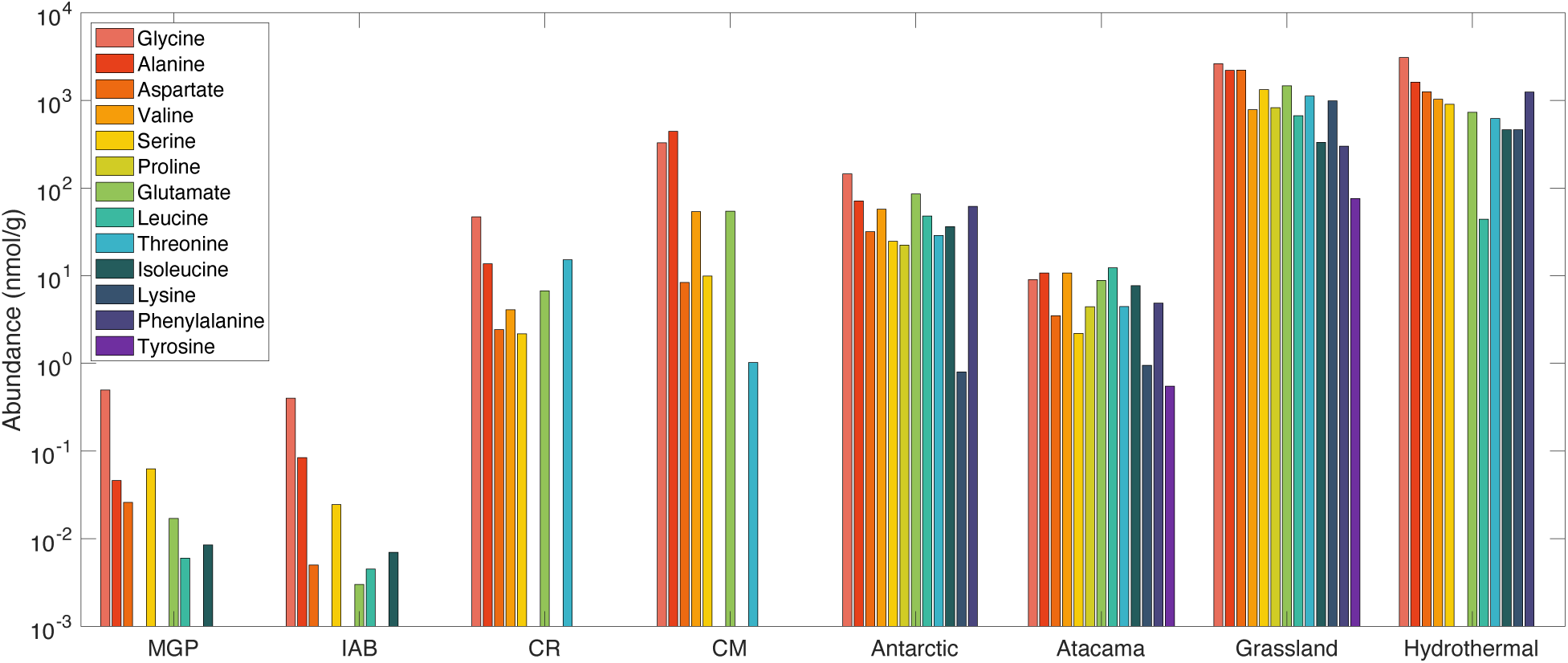
Distribution patterns of meteoritic ⍺-amino acids in different meteorite groups and amino acids in various biogenic samples. Abundance data was averaged following the concentrations reported in tables S1 and S2. The amino acids are placed in the consensus evolutionary order predicted by Trifonov 2009. Meteorite group key: MGP (Main Group Pallasites), IAB (Iron Meteorites Group AB), CR (Renazzo-type chondrites), and CM (Mighei- type chondrites).

**Figure S7.**
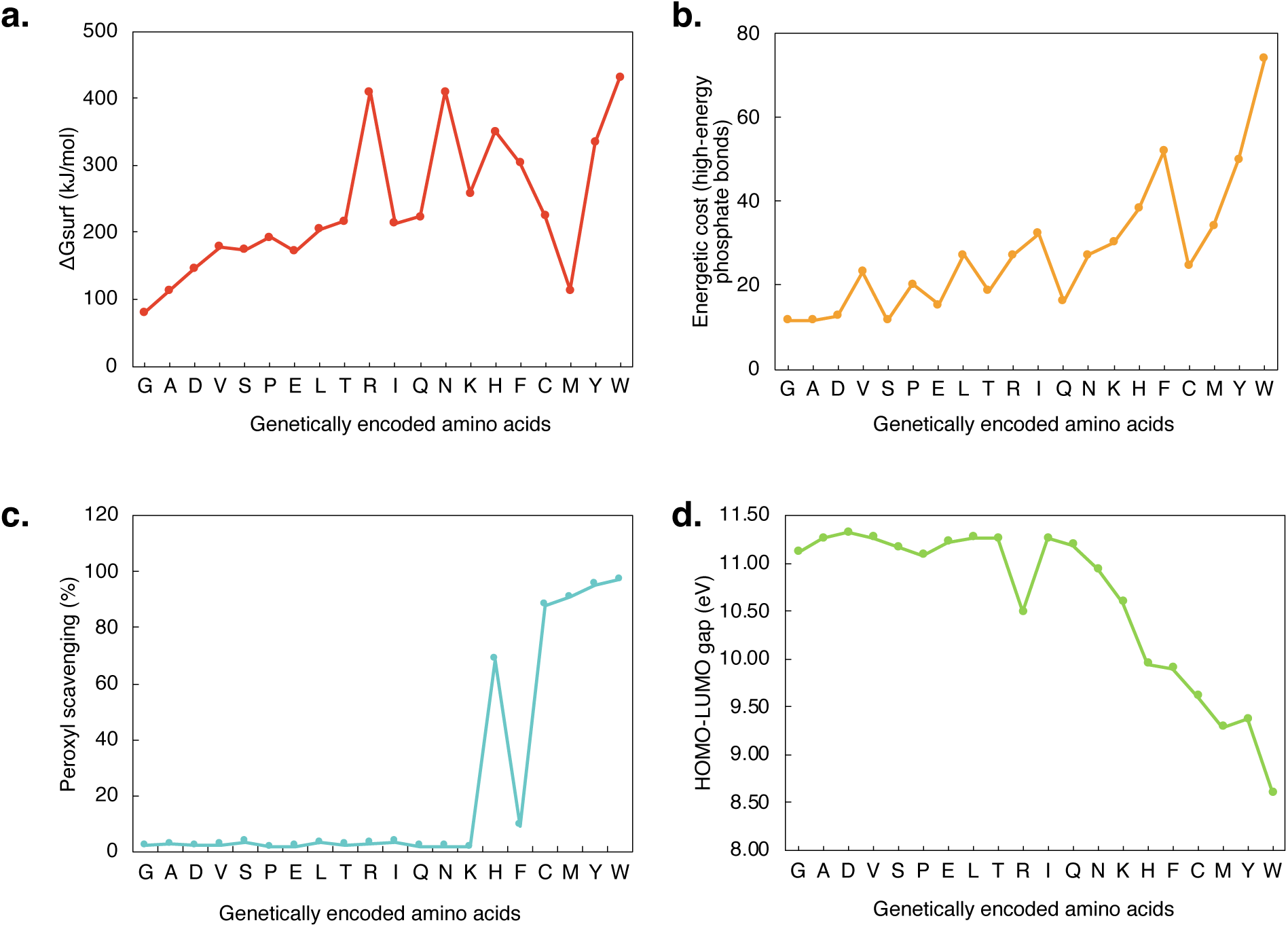
Patterns of various quantum, redox, and thermodynamic properties of amino acids. Consensus order of appearance of proteinogenic amino acids within the genetic code, as predicted by Trifonov 2009, in function of their: **a,** Gibbs free energies of net amino acid synthesis reactions in surface seawater (as reported by Amend and Shock 1998); **b,** energetic cost of biosynthesis in *E. coli* and *B. subtilis* (as reported by Akashi and Gojobori 2002); **c,** reactivity towards peroxyl radicals and **d,** HOMO-LUMO gap values (as reported by Granold *et al*. 2018). Refer to Dataset 3 for quantitative data regarding each property.

**Figure S8.**
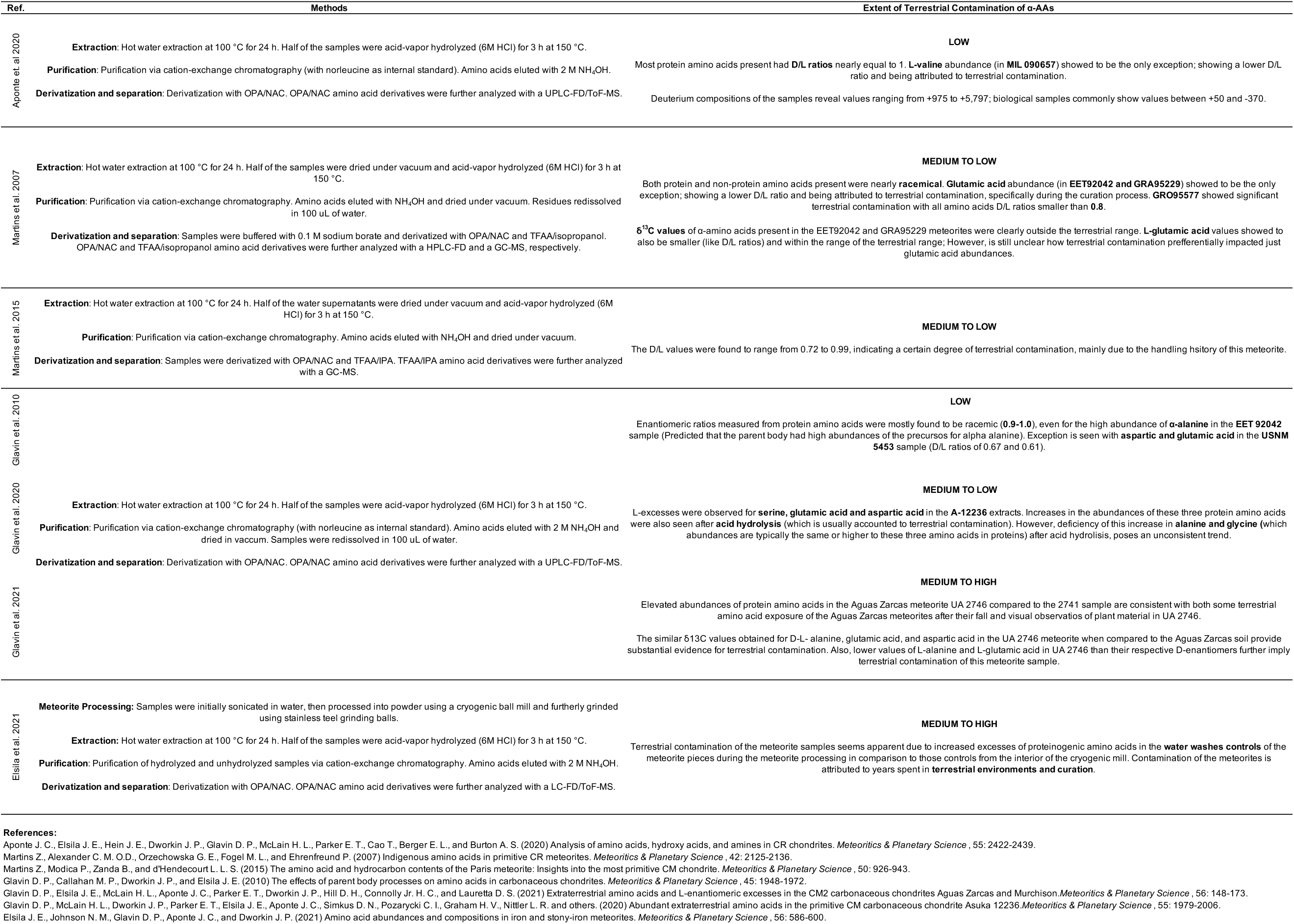
Methods employed in the extraction of amino acids from meteroites studied, along the degree of terrestrial contamination of the samples.

**Figure S9.**
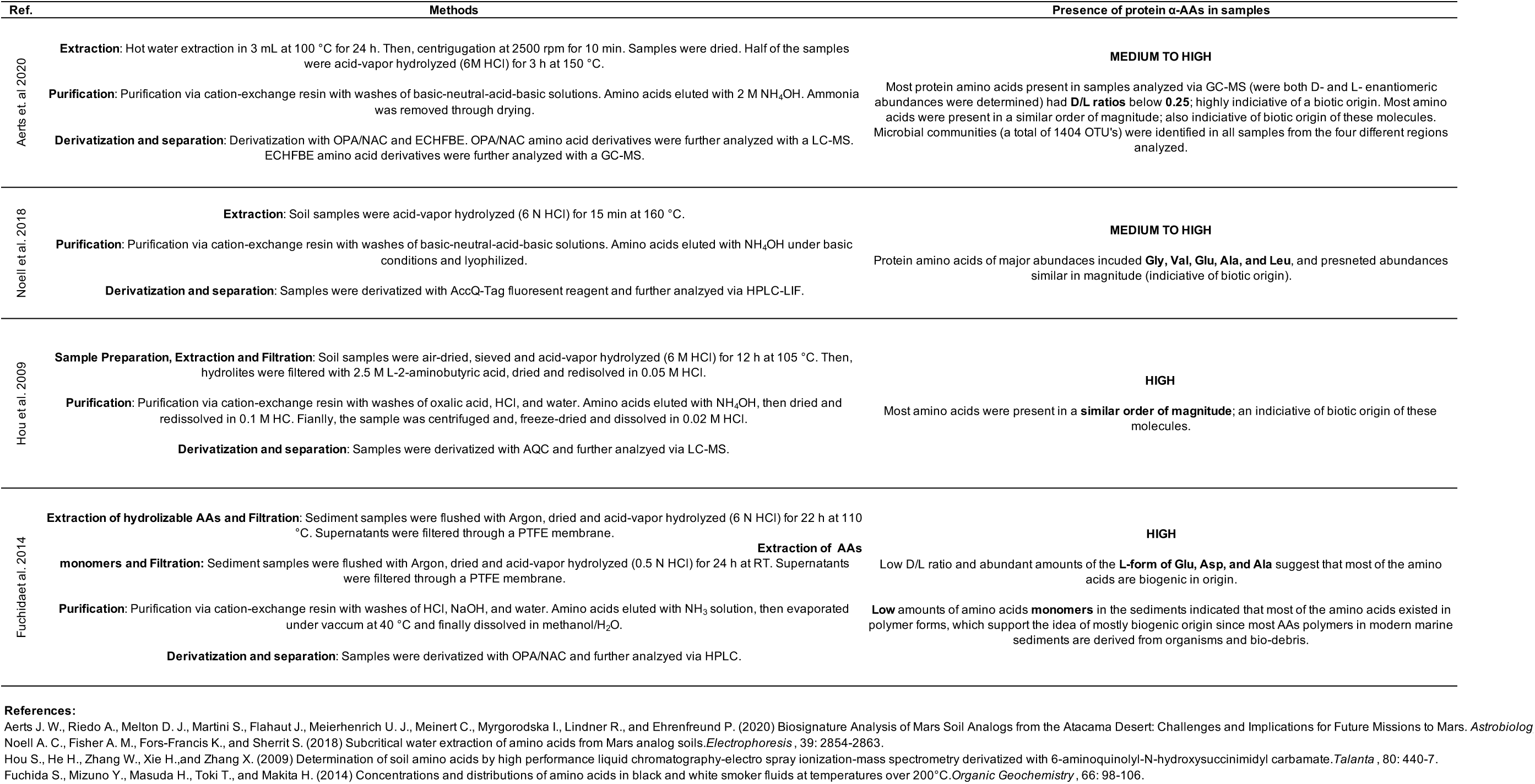
Methods employed in the extraction of amino acids from environemental samples studied, along the presence of protein α-AAs on each.

**Table S1.**
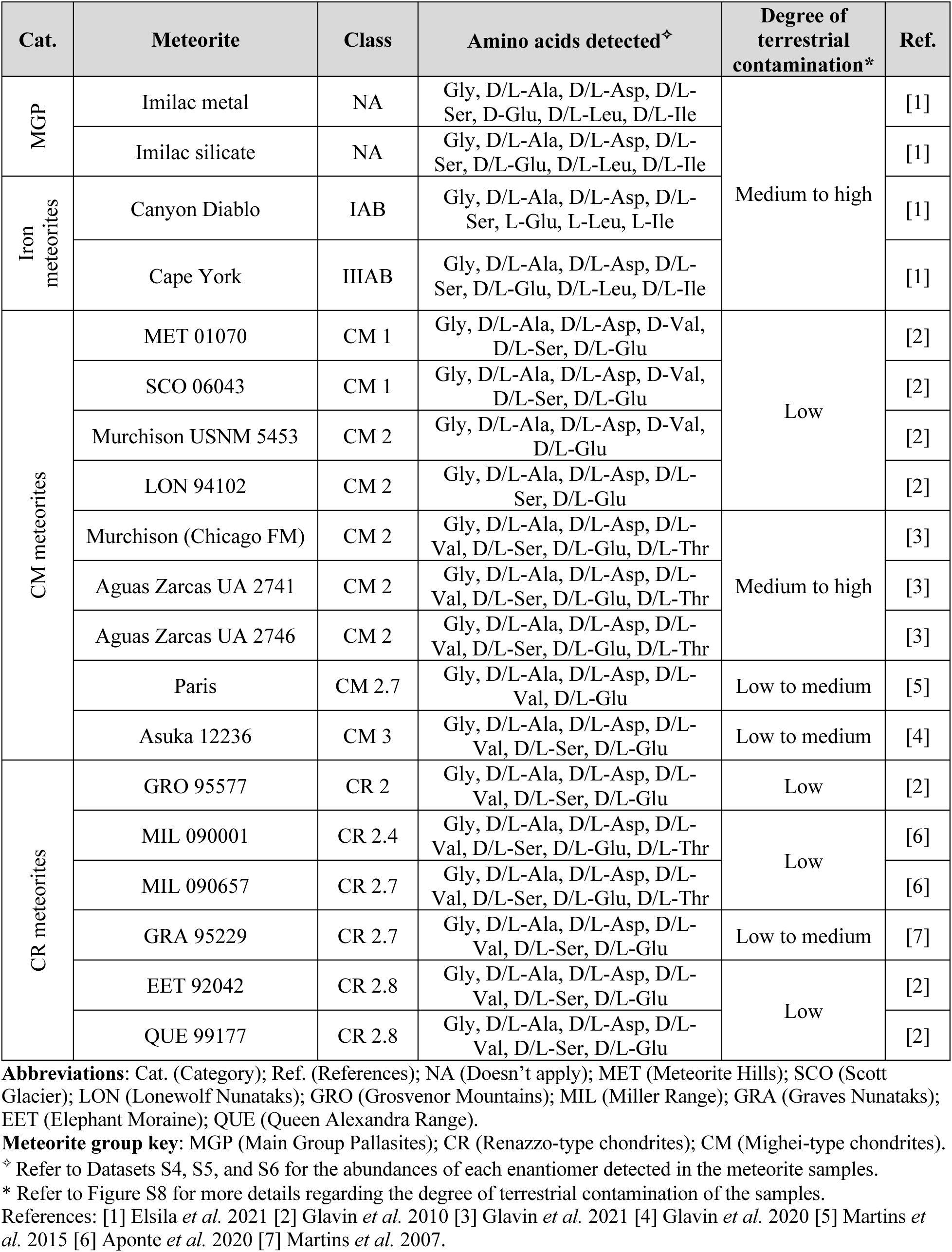
Meteorites samples studied with their respective classification.

| Cat. | Meteorite | Class | Amino acids detected<sup>‡</sup> | Degree of terrestrial contamination* | Ref. |
| --- | --- | --- | --- | --- | --- |
| MGP | Imilac metal | NA | Gly, D/L-Ala, D/L-Asp, D/L-Ser, D-Glu, D/L-Leu, D/L-Ile | Medium to high | [1] |
|  | Imilac silicate | NA | Gly, D/L-Ala, D/L-Asp, D/L-Ser, D/L-Glu, D/L-Leu, D/L-Ile |  | [1] |
| Iron meteorites | Canyon Diablo | IAB | Gly, D/L-Ala, D/L-Asp, D/L-Ser, L-Glu, L-Leu, L-Ile |  | [1] |
|  | Cape York | IIIAB | Gly, D/L-Ala, D/L-Asp, D/L-Ser, D/L-Glu, D/L-Leu, D/L-Ile |  | [1] |
| CM meteorites | MET 01070 | CM 1 | Gly, D/L-Ala, D/L-Asp, D-Val, D/L-Ser, D/L-Glu | Low | [2] |
|  | SCO 06043 | CM 1 | Gly, D/L-Ala, D/L-Asp, D-Val, D/L-Ser, D/L-Glu |  | [2] |
|  | Murchison USNM 5453 | CM 2 | Gly, D/L-Ala, D/L-Asp, D-Val, D/L-Glu |  | [2] |
|  | LON 94102 | CM 2 | Gly, D/L-Ala, D/L-Asp, D/L-Ser, D/L-Glu |  | [2] |
|  | Murchison (Chicago FM) | CM 2 | Gly, D/L-Ala, D/L-Asp, D/L-Val, D/L-Ser, D/L-Glu, D/L-Thr | Medium to high | [3] |
|  | Aguas Zarcas UA 2741 | CM 2 | Gly, D/L-Ala, D/L-Asp, D/L-Val, D/L-Ser, D/L-Glu, D/L-Thr |  | [3] |
|  | Aguas Zarcas UA 2746 | CM 2 | Gly, D/L-Ala, D/L-Asp, D/L-Val, D/L-Ser, D/L-Glu, D/L-Thr |  | [3] |
|  | Paris | CM 2.7 | Gly, D/L-Ala, D/L-Asp, D/L-Val, D/L-Glu | Low to medium | [5] |
|  | Asuka 12236 | CM 3 | Gly, D/L-Ala, D/L-Asp, D/L-Val, D/L-Ser, D/L-Glu | Low to medium | [4] |
| CR meteorites | GRO 95577 | CR 2 | Gly, D/L-Ala, D/L-Asp, D/L-Val, D/L-Ser, D/L-Glu | Low | [2] |
|  | MIL 090001 | CR 2.4 | Gly, D/L-Ala, D/L-Asp, D/L-Val, D/L-Ser, D/L-Glu, D/L-Thr | Low | [6] |
|  | MIL 090657 | CR 2.7 | Gly, D/L-Ala, D/L-Asp, D/L-Val, D/L-Ser, D/L-Glu, D/L-Thr |  | [6] |
|  | GRA 95229 | CR 2.7 | Gly, D/L-Ala, D/L-Asp, D/L-Val, D/L-Ser, D/L-Glu | Low to medium | [7] |
|  | EET 92042 | CR 2.8 | Gly, D/L-Ala, D/L-Asp, D/L-Val, D/L-Ser, D/L-Glu | Low | [2] |
|  | QUE 99177 | CR 2.8 | Gly, D/L-Ala, D/L-Asp, D/L-Val, D/L-Ser, D/L-Glu |  | [2] |

**Abbreviations:** Cat. (Category); Ref. (References); NA (Doesn't apply); MET (Meteorite Hills); SCO (Scott Glacier); LON (Lonewolf Nunataks); GRO (Grosvenor Mountains); MIL (Miller Range); GRA (Graves Nunataks); EET (Elephant Moraine); QUE (Queen Alexandra Range).
**Meteorite group key:** MGP (Main Group Pallasites); CR (Renazzo-type chondrites); CM (Mighei-type chondrites).
<sup>‡</sup> Refer to Datasets S4, S5, and S6 for the abundances of each enantiomer detected in the meteorite samples.
\* Refer to Figure S8 for more details regarding the degree of terrestrial contamination of the samples.
References: [1] Elsila *et al.* 2021 [2] Glavin *et al.* 2010 [3] Glavin *et al.* 2021 [4] Glavin *et al.* 2020 [5] Martins *et al.* 2015 [6] Aponte *et al.* 2020 [7] Martins *et al.* 2007.

**Table S2.** Environmental samples studied with their respective collection site.

| Cat. | Sample ID | Location | Amino acids detected<sup>†</sup> | Degree of biogenic origin* | Ref. |
| --- | --- | --- | --- | --- | --- |
| Atacama | ATA03 | Salar Laco (margin) | Gly, L-Ala, L-Asp, L-Val, L-Ser, L-Glu, L-Leu, L-Thr, L-Ile, L-Phe | Medium to high | [1] |
|  | ATA05 | Laguna Tuyajto (margin) | Gly, L-Ala, L-Asp, L-Val, L-Ser, L-Glu, L-Leu, L-Thr, L-Ile, L-Phe |  | [1] |
|  | ATA12 | Salar Aguas Calientes 3 (margin) | Gly, D/L-Ala, L-Asp, L-Val, L-Ser, D/L-Pro, D/L-Glu, D/L-Leu, D/L-Ile, D/L-Phe |  | [1] |
|  | ATA15 | Salar de Atacama Peine Road (margin) | Gly, D/L-Ala, L-Asp, L-Val, D/L-Pro, L-Glu, D/L-Leu, D/L-Ile, D/L-Phe |  | [1] |
|  | AT45-A3 | Yungay Hill | Gly, D/L-Ala, D/L-Asp, L-Val, D/L-Ser, D/L-Pro, D/L-Glu, D/L-Leu, D/L-Thr, D/L-Ile, D/L-Phe, D/L-Lys, D/L-Tyr | Medium to high | [2] |
| Antarctica |  | Wright Valley | Gly, D/L-Ala, D/L-Asp, L-Val, D/L-Ser, D/L-Pro, D/L-Glu, D/L-Leu, D/L-Thr, D/L-Ile, D/L-Phe, D/L-Lys, D/L-Tyr |  | [2] |
| Grassland | Mollisol-1 | Hailun, Heilongjian, China | Gly, D/L-Ala, D/L-Asp, L-Val, D/L-Ser, D/L-Pro, D/L-Glu, D/L-Leu, D/L-Thr, D/L-Ile, D/L-Phe, D/L-Lys, D/L-Tyr | High | [3] |
| Hydrothermal Systems | PC-1 | Izena Cauldron | Gly, D/L-Ala, D/L-Asp, L-Val, D/L-Ser, D/L-Glu, D/L-Leu, D/L-Thr, D/L-Ile, D/L-Phe, D/L-Lys | High | [4] |
|  | PC-3 | Yoron Cauldron | Gly, D/L-Ala, D/L-Asp, L-Val, D/L-Ser, D/L-Glu, D/L-Leu, D/L-Thr, D/L-Ile, D/L-Phe, D/L-Lys |  | [4] |

Abbreviations: Cat. (Category), Ref. (References).
<sup>†</sup> Refer to Dataset S7 for the abundances of each enantiomer detected in the environmental samples.
\* Refer to Figure S9 for more details regarding the degree of biogenic origin of the amino acids in each sample.
References: [1] Aerts *et al.* 2020 [2] Noell *et al.* 2018 [3] Hou *et al.* 2009 [4] Fuchida *et al.* 2014.

**Table S3.**
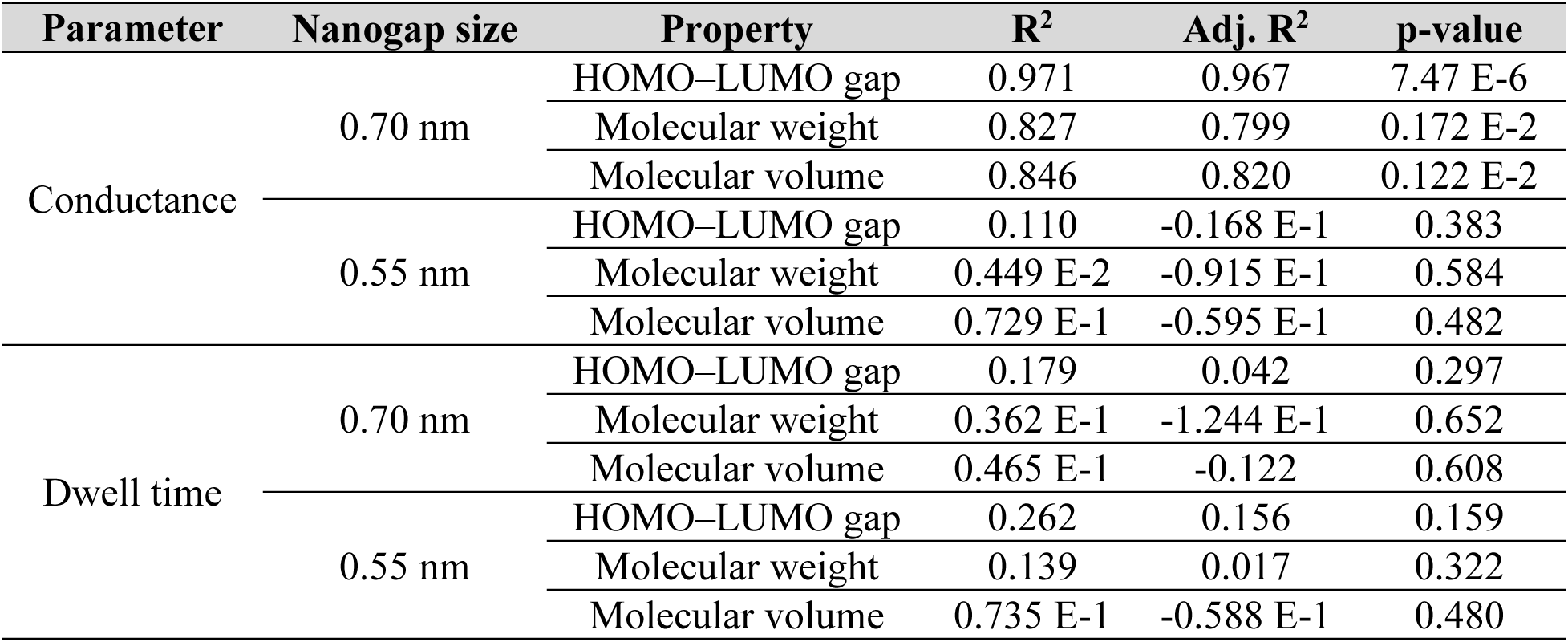
Statistical summary of the linear regression plots. Fitted by plotting the conductance and dwell time measurements of Ohshiro *et al*. 2014 for 12 amino acids using 0.7- nm- and 0.55-nm-nanogap electrodes against the specific amino acid’s molecular weight, volume and HOMO–LUMO gap.

---

**Dataset S1.** Single-molecule conductance and duration time for 13 amino acid molecules using 0.7-nm- and 0.55-nm-nanogap electrodes.

| Amino Acid | Abbreviation | Symbol | Gap_nm | Conductance_pS | Spread_pS | Dwell_time_ms | Spread_ms |
| --- | --- | --- | --- | --- | --- | --- | --- |
| Aspartic Acid | Asp | D | 0.7 | 183 | 22 | 0.98 | 0.63 |
| Isoleucine | Ile | I | 0.7 | 128 | 20 | 1.15 | 0.42 |
| Glutamic Acid | Glu | E | 0.7 | 168 | 34 | 1.57 | 0.46 |
| Proline | Pro | P | 0.7 | 235 | 40 | 2.54 | 0.59 |
| Histidine | His | H | 0.7 | 609 | 55 | 3.46 | 0.8 |
| Phenylalanine | Phe | F | 0.7 | 690 | 64 | 1.53 | 0.49 |
| Tyrosine | Tyr | Y | 0.7 | 874 | 84 | 1.54 | 0.47 |
| Tryptophan | Trp | W | 0.7 | 953 | 130 | 2.52 | 1.05 |
| Aspartic Acid | Asp | D | 0.55 | 200 | 45 | 0.95 | 0.43 |
| Isoleucine | Ile | I | 0.55 | 161 | 18 | 1.09 | 0.47 |
| Glutamic Acid | Glu | E | 0.55 | 234 | 82 | 0.6 | 0.59 |
| Proline | Pro | P | 0.55 | 670 | 31 | 2.13 | 1.1 |
| Histidine | His | H | 0.55 | 1088 | 133 | 1.53 | 0.5 |
| Lysine | Lys | K | 0.55 | 548 | 106 | 3.54 | 0.87 |
| Cysteine | Cys | C | 0.55 | 431 | 55 | 6.5 | 3.2 |
| Methionine | Met | M | 0.55 | 266 | 40 | 1.4 | 1.0 |
| Leucine | Leu | L | 0.55 | 332 | 74 | 1.2 | 0.5 |

**Dataset S2.**
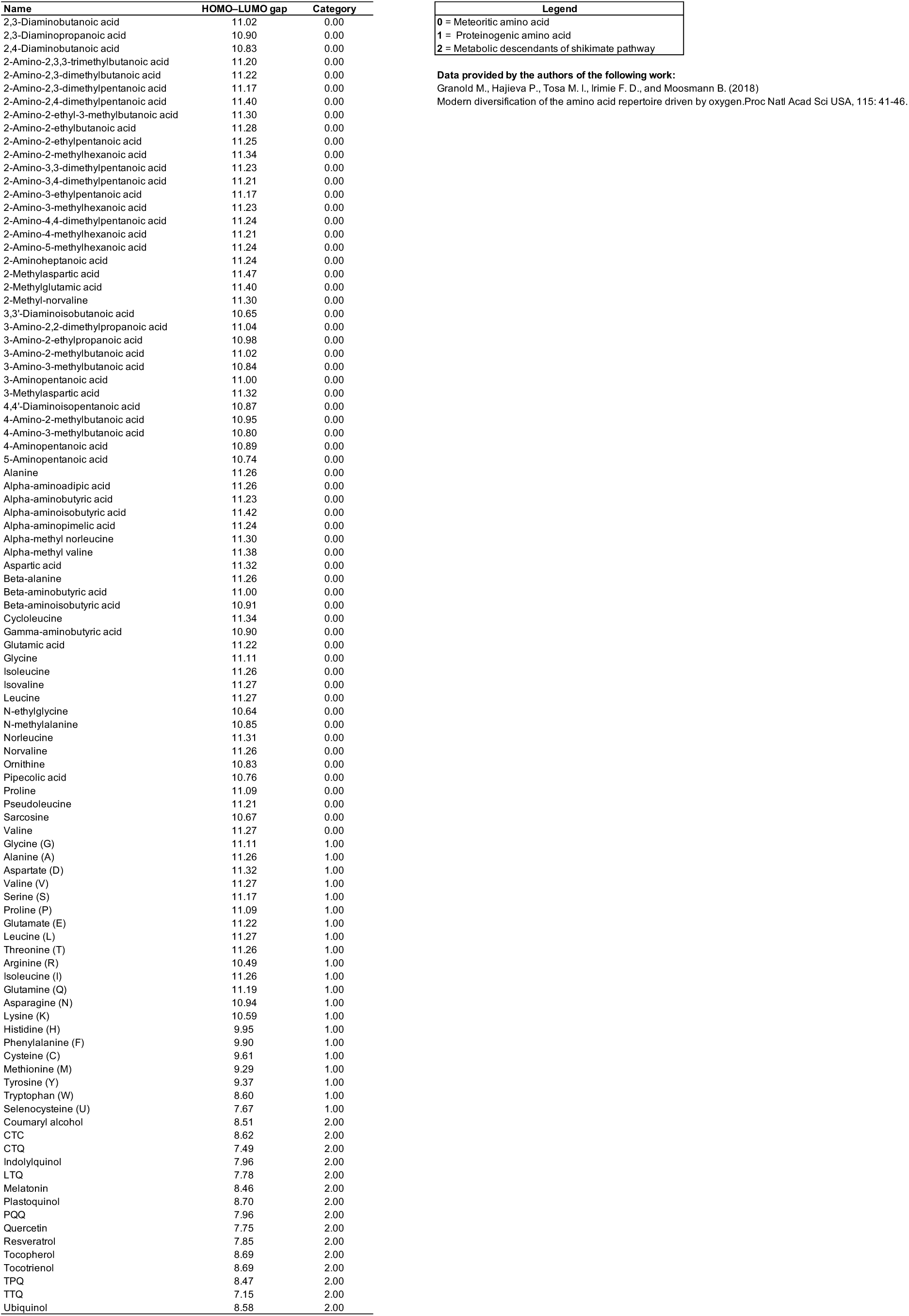
HOMO–LUMO gaps of 62 Murchison meteorite AAs, 21 genetically encoded AAs, and various metabolic descendants of the shikimate pathway calculated using semiempirical methods (AM1).

**Dataset S3.**
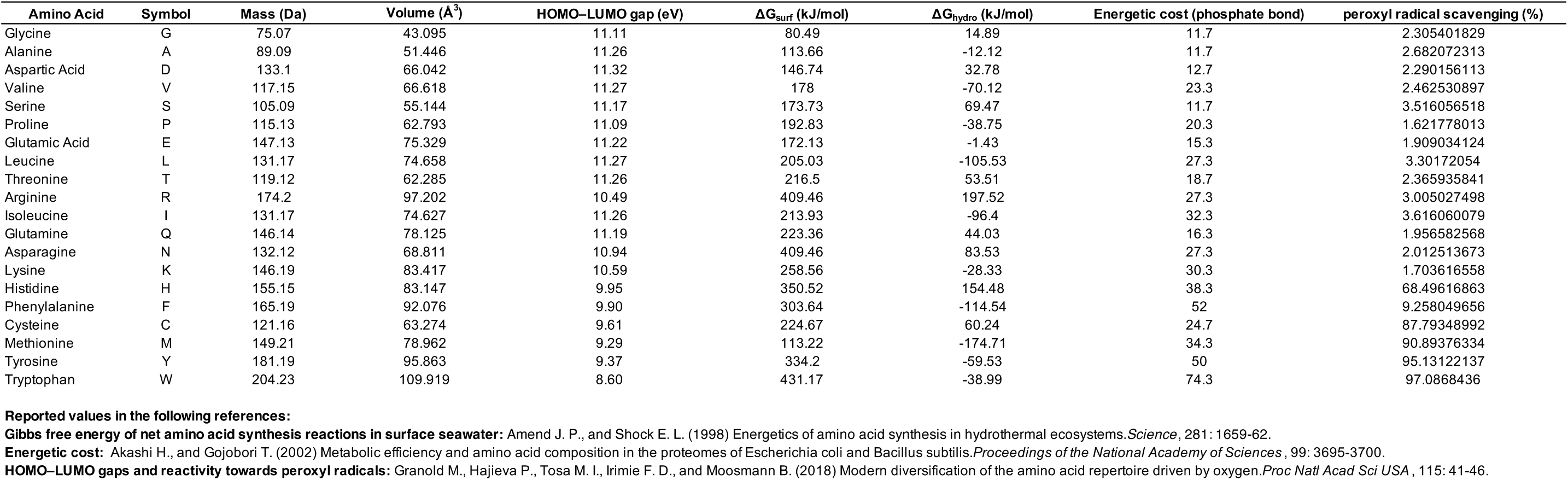
Quantitative data of studied quantum, redox, and thermodynamic properties of amino acids.

**Dataset S4.** Amino acid abundances (nmol/g) in meteorites samples of the Main Group Pallasites and Iron meteorites group AB.

| Amino acid | MGP |  |  |  | IAB |  |  |  |
| --- | --- | --- | --- | --- | --- | --- | --- | --- |
|  | Elsila et al. 2021 (nmol/g) |  |  |  |  |  |  |  |
|  | Imilac metal | SD | Imilac silicate | SD | Canyon Diablo | SD | Cape York | SD |
| D-aspartic acid | 0.0013 | SM | 0.0062 | SM | Below limit | NA | 0.0019 | SM |
| L-aspartic acid | Below limit | NA | 0.0446 | SM | 0.0035 | 0 | 0.0045 | SM |
| D-glutamic acid | Below limit | NA | 0.0016 | SM | Not detected | NA | Below limit | SM |
| L-glutamic acid | Not detected | NA | 0.0156 | SM | 0.0052 | 0 | 0.0013 | SM |
| D-serine | 0.002 | SM | 0.0146 | SM | 0.0017 | 0 | 0.0015 | SM |
| L-serine | 0.0035 | SM | 0.1044 | SM | 0.0387 | 0 | 0.0078 | SM |
| glycine | 0.1902 | SM | 0.8024 | SM | 0.5274 | 0.06 | 0.2728 | SM |
| D-alanine | 0.0069 | SM | 0.0164 | SM | 0.0208 | 0 | 0.0078 | SM |
| L-alanine | 0.0107 | SM | 0.0573 | SM | 0.1238 | 0.02 | 0.015 | SM |
| D-isoleucine | Below limit | NA | Below limit | NA | Not detected | NA | Below limit | NA |
| L-isoleucine | 0.0018 | SM | 0.0149 | SM | 0.0131 | 0 | 0.0014 | SM |
| D-leucine | Below limit | NA | Below limit | NA | Not detected | NA | Below limit | NA |
| L-leucine | 0.0016 | SM | 0.0096 | SM | 0.0081 | 0 | 0.0009 | SM |

SM: Single Measurement
NA: Doesn't apply

**Dataset S5.**
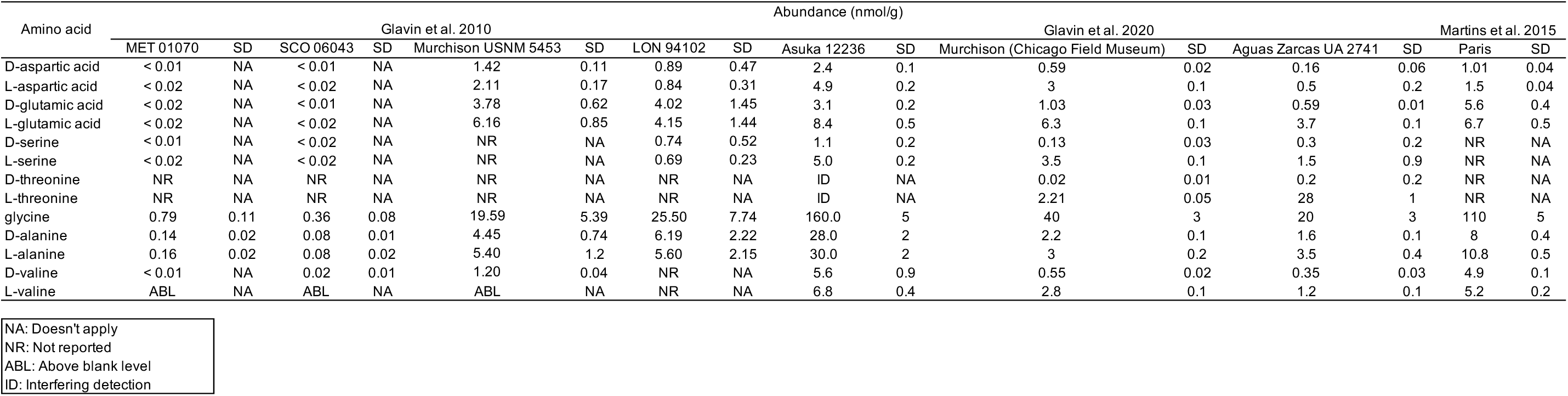
Amino acid abundances (nmol/g) in meteroites samples of the Mighei-type chondrites group.

**Dataset S6.**
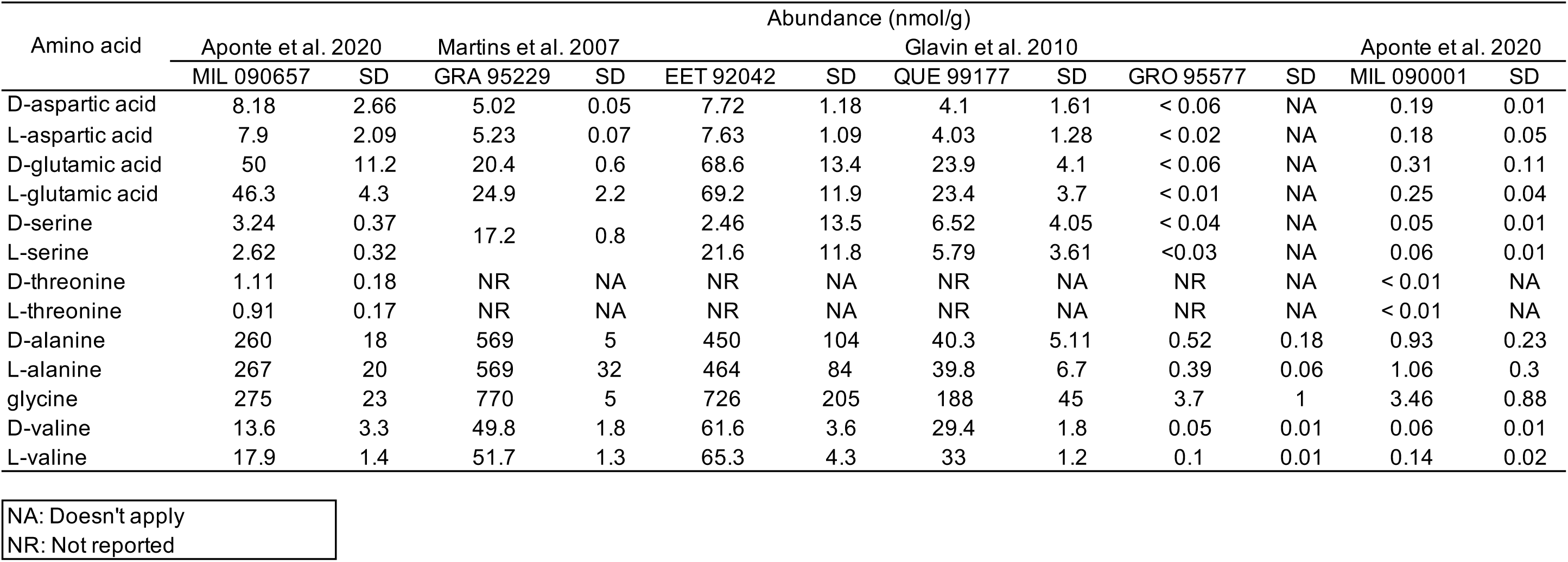
Amino acid abundances (nmol/g) in meteroite samples of the Renazzo-type chondrites group.

**Dataset S7.**
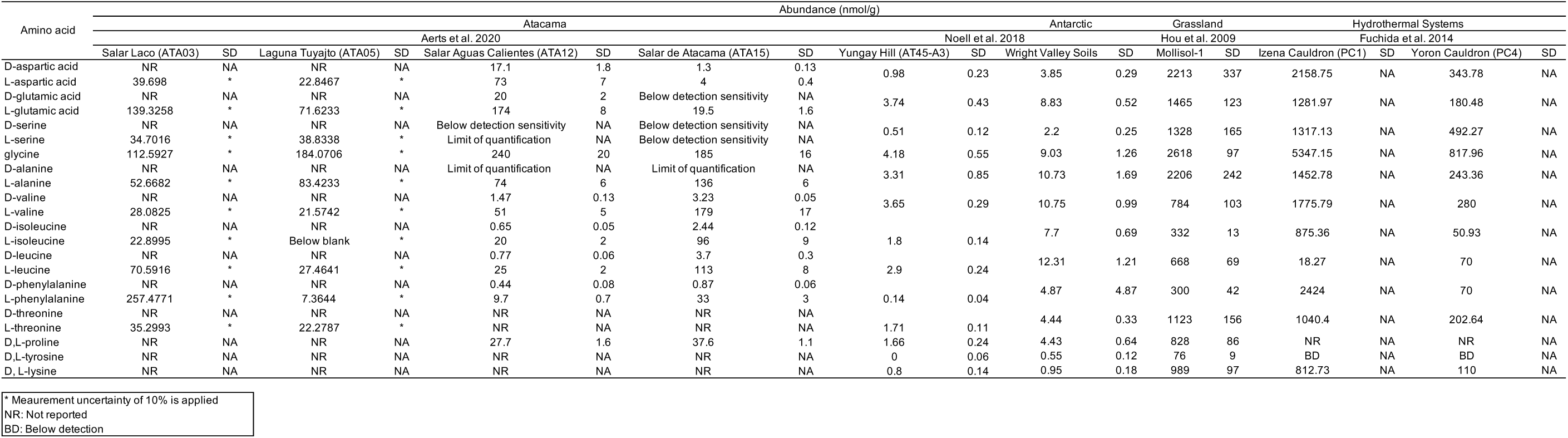
Amino acid abundances (nmol/g) in environmental samples from the Atacama, Antarctica, grassland, and hydrothermal systems.

## Notes

### Competing Interest Statement

The authors have declared no competing interest.

